# In depth characterization of an archaeal virus-host system reveals numerous virus exclusion mechanisms

**DOI:** 10.1101/2022.10.18.512658

**Authors:** Coraline Mercier, Daniela Thies, Ling Zhong, Mark J. Raftery, Ricardo Cavicchioli, Susanne Erdmann

**Affiliations:** Max Planck Institute for Marine Microbiology, Archaeal Virology, Celsiusstrasse 1, 28359, Bremen, Germany; Bioanalytical Mass Spectrometry Facility, The University of New South Wales, Sydney, New South Wales, 2052, Australia; School of Biotechnology and Biomolecular Sciences, The University of New South Wales, Sydney, New South Wales, 2052, Australia

## Abstract

Archaeal head-tailed viruses appear, at first sight, very closely related to head-tailed bacteriophages simply due to morphological similarities and similar life cycles. However, they encounter host cells that are very different from bacteria and share characteristics, that greatly influence virus life cycles, with eukaryotes. Here we present an in-depth characterization of the archaeal head-tailed virus, HRTV-Dl1, isolated from Deep Lake, Antarctica. The host *Halorubrum lacusprofundi* exhibits a large arsenal of virus exclusion mechanisms, indicating a long ongoing arms race with viruses. However, we uncover that the majority of this arsenal was lost spontaneously in a strain grown under non-challenging laboratory conditions. By challenging both the parental strain and the sensitive strain with HRTV-DL1, we discovered a number of putative virus exclusion mechanisms that are only activated in the sensitive strain upon the lack of defense systems present in the parental strain. We identify virus exclusion mechanisms that are also common in bacteria, mechanisms that are unique to archaea, and a potential mechanisms involving the archaeal homolog of the eukaryotic ORC1 and CDC6. We identify one of two S-layer proteins as primary receptor for HRTV-DL1, demonstrating that the presence of two different S-layer proteins in one strain provides a strong advantage in the arms race with viruses. Our results clearly reflect the differences between bacterial and archaeal head-tailed viruses. Finally, we observed that our intention to isolate a clean and stable model virus-host system led to the generation of a virus-host pair with reduced genomes. This model system is great to study in the laboratory, but barely reflects the entire spectrum of virus-host interactions as they would occur in the environment, emphasizing the importance of combining wet lab data with environmental data.

## Introduction

Deep Lake is a cold, marine-derived hypersaline lake in Antarctica with temperatures ranging between −20°C in the bulk of the lake and up to +12°C at the surface [1]. The lake is globally isolated and is characterized by a very limited microbial diversity [2]. The community is dominated by four halophilic archaeal species (haloarchaea) accounting for ~72% of the population, with *Hst. litchfieldiae* tADL accounting for 43.5%, halophilic archaeon DL31 for 18.2%, *Halorubrum lacusprofundi* for 9.9% and *Halobacterium* DL1 for 0.9% [3]. These species, except one (*Halohasta litchfieldae* tADL), exhibit one main chromosome and one or several secondary replicons (secondary chromosomes or megaplasmids). The secondary replicons, that are also common in other haloarchaea [4], were shown to be extremely diverse even between strains of the same species, in particular in *Hrr. lacusprofundi* [5]. While some of these secondary replicons strongly interact with the main chromosomes and exhibit some genes that are essential [6], others have been lost and shown to be non-essential [7]. Nevertheless, the majority of genes encoded on secondary replicons are thought to allow adaptation to changing environmental conditions, but are possibly not essential under stable laboratory conditions. *Hrr. lacusprofundi* exhibts three replicons, the main chromosome CHR1, a secondary chromosome CHR2 and the megaplasmid pHLAC01. One ribosomal rRNA operon is located on CHR2 and a number of virus defense systems, including two CRISPR-Cas systems, are found on both CHR2 and pHLAC01. A metagenomics study on Antarctic salt lakes revealed that the main chromosome of *Hrr. lacusprofundi* is highly conserved between species, but the secondary chromosomes and megaplasmids are very diverse and responsible for the genetic diversity of the species [5]. A plasmid mobilized between *Hrr. lacusprofundi* strains in specialized membrane vesicles was proposed to be involved in generating this diversity [8].

Owing to its low-complexity community and the closed nature of the system, Deep Lake is ideal for studying virus-host interactions and the impact of viruses on a microbial community. Only one putative primary producer, the green alga *Dunaliella* sp., was identified in the lake, however, no grazing organisms were detected. Therefore, while viruses are important regulators of community structure in all environments [9], we suggest that they play a particular important role in Deep Lake as one of the major players influencing microbial community structure. Several omics-based studies on Deep Lake revealed an ongoing arms race between the Deep Lake haloarchaea and viruses, and suggest that viruses are one of the drivers of genetic exchange in the community [3, 5, 10]. However, in depth characterization of virus-host interactions, including the life cycle of viruses, the host range and the influence on host viability, is only feasible on an isolated virus-host system. Even though the major microbial players have been isolated, no virus was isolated from Deep Lake so far. In an effort to understand the influence of viruses on Deep Lake haloarchaea we envisaged to isolate and characterize representative virus-host systems, subsequently allowing the assessment of the respective relationships in their natural environment using metagenomics data from several years and different seasons.

While studies on archaeal viruses in general are still significantly underrepresented, the majority of isolated viruses to date have been described for members of Halobacteriota, Methanobacteriota and Thermoproteota phyla. Haloarchaea (belonging to the Halobacteriota) are infected by diverse viruses including head-tailed viruses, pleomorphic viruses, spherical viruses and spindle shaped viruses [11]. A number of haloarchaeal viruses exhibit lysogenic and chronic life cycles, however, the majority of isolated viruses exhibit lytic life cycles, reflecting a methodical bias caused by using plaque assays for isolation, selecting mostly for lytic viruses [12]. A recent study described a number of new haloarchaeal virus genomes and proposed a new classification of archaeal tailed viruses (arTV) [13]. Even though arTV appear morphological similar to head-tailed bacteriophages, they encounter cells that are dramatically different, in particular with respect to the cell surface, replication, transcription and translation mechanisms. Nevertheless, to date only very few arTV infecting haloarchaea have been characterized in more depth [11, 13].

Recent studies, using bioinformatics approaches, identified an enormous diversity of previously unrecognised bacterial and archaeal virus exclusion mechanisms. While some of these defence systems have been characterized in bacteria [14], their activity in archaea remains to be elucidatedAdditionally, archaeal viruses are in general significantly more diverse than isolated bacterial viruses, raising the question as to which extent the currently known defence systems are active against which range of archaeal viruses. Moreover, the fact that archaea share characteristics with eukaryotes potentially expands the range of potential virus exclusion mechanisms, that are not present in bacteria, to be discovered.

Here we present an in-depth characterisation of an isolated virus-host system from Deep Lake including the characterisation of the virus life cycle, determination of the virus host range, analysis of the host transcriptional response to virus infection and the characterisation of virus escape mutants. We uncover the genomic plasticity of both the host and the virus. We determine the host receptor for the virus and discover a number of new potential virus exclusion mechanisms. Establishing this isolated virus-host system indeed allows to uncover many molecular aspects of virus–host interactions that are not extractable from ‘omics’ data. However, we also realized that the isolated system only allows a glimpse into the possible interactions ongoing in the natural environment, which perhaps can be elucidated using ‘omics’ data.

## Results and Discussion

### Isolation of a lytic virus that infects Halorubrum lacusprofundi ACAM34 (DSM 5036)

A virus with head-tailed morphology (Figure 1b) was obtained from the culture supernatant of a *Halorubrum lacusprofundi* strain, that was isolated from a sample taken in 2014 from Deep Lake, Antarctica [1]. Analytic restriction digest of DNA isolated from the viral lysate revealed a dsDNA genome of approximately 60 kb (Figure 1a). The lysate produced plaques on *Hrr. lacusprofundi* strain ACAM34 [15] that was isolated from a Deep Lake sample in 1988, while it remains elusive when the sample was actually obtained from Deep Lake. Lysis was also observed in liquid cultures of this strain. The viral lysate was formerly describe as DLHTHV [5], however, we rename it hereby as HRTV-DL (**H**alo**R**ubrum **T**ailed **V**irus-**D**eep **L**ake), according to the nomenclature that was previously used for archaeal tailed viruses (arTV) [13]. HRTV-DL is the first virus infecting *Hrr. lacusprofundi* that has been isolated from Deep Lake and, interestingly, even though the virus was isolated at least 26 years later than the host, it is still able to generate a successful infection. *Hrr. lacusprofundi* ACAM34 was used to purify the virus through several rounds of plaque assays. When analysing genomic DNA of virus particles propagated from single plaques, we observed a remarkable high genomic variability (Figure S1). Such a genomic variation has previously been observed for *Halobacterium salinarium* virus ØH [16]. After two rounds of plaque purification, we isolated a variant that appeared to have a stable genome through several series of infection (P2V1, Figure S1c) and we choose this variant, HRTV-DL1, for further characterization. Virus particles analysed by electron microscopy exhibit a head-tailed morphology, with a head diameter of about 50 nm and a non-contractile tail of approximately 80 nm-100 nm in length (Figure 1b, Figure S2).

**Figure 1.**
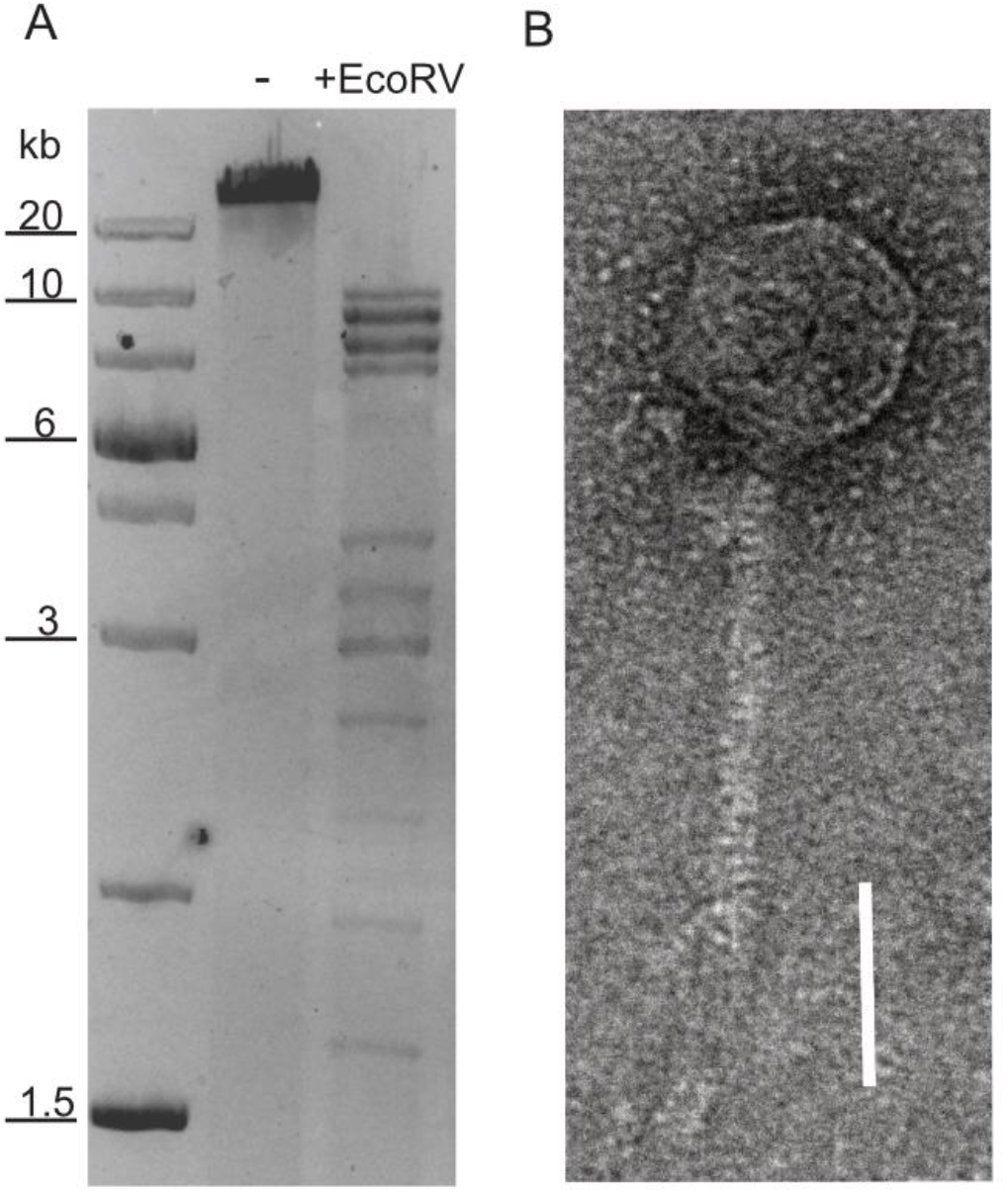
Nucleic acid content and particle of HRTV-DL. **(A)**Total DNA of HRTV-DL undigested (-) and digested with *EcoRV* (+EcoRV). MW size marker is shown to the left of the gel (GeneRuler 1 kb Plus DNA Ladder, Thermo Fisher Scientific). DNA was separated on 1% agarose gels and stained with SYBR™ Safe DNA stain. **(B)**Electron micrograph of HRTV-DL particle. Sample was negatively stained with 2% uranyl acetate. Size bar: 50nm. Original images have been modified by cropping to improve visual presentation.

### Genome comparison of the original virus lysate and the isolated variant genome reveals genome plasticity of the virus and unintentional selection for a lytic virus variant

DNA isolated from HTRV-DL1 particles appeared sensitive to Exonuclease treatment, while the majority of virus DNA isolated from infected cells was shown to be circular (Figure S3), indicative of circularisation of the viral genome after injection into the host. Analytic digest of virus and host DNA with *Dpn*I, showed that both the host and the virus genome are dam methylated. Methylation of the virus genome likely supports successful proliferation of the virus within its host, allowing the virus to escape host Restriction-modification (R-M) defence systems. Both the original lysate (HRTV-DL) and the isolated variant HRTV-DL1 were subjected to genome sequencing. The genome of HRTV-DL assembled in a number of contigs, that could not be computationally connected, and the genome of HRTV-DL1 assembled into a single circular contig of 37.7 kb. Since the HRTV-DL1 genome was shown to be linear within the virus particle but assembles as a circular contig, with no indication in the sequence data for discrete ends, we conclude that the genome is circularly permuted and terminally redundant. Contigs of HRTV-DL could manually be connected by PCR into one contig of 66.2kb, similar to what has been estimated by analytic restriction digest (Figure 1a). We assume that we were able to close it to only one contig, because the genome is circularly permuted and terminally redundant, however, it would never appear as one complete genome in one virus particle. Comparison of the two genomes revealed that HRTV-DL contains several duplicated regions and a few regions that are unique (Figure S4). This included 10 hypothetical proteins, 2 integrases and one adenine-specific DNA methylase. Additionally, the phage tail tape measure protein is 28aa shorter in HRTV-DL1 (Table S1). We suggest that upon initial isolation of HRTV-DL, the viral lysate contained a mixture of different virus variants that shared the majority of their genome. Co-infection allowed the recombination of the different genomic regions and the generation of diverse virus variants as observed during plaque purification (Figure S1). Interestingly, none of the integrases that are present in the HRTV-DL lysate was found in the isolated variant HRTV-DL1 that exclusively exhibits a lytic cycle. By using plaque assay for purification, we selected for a virus variant with a lytic cycle and thereby lost variants that would be able to integrate into the host genome and exhibit a lysogenic life cycle. After obtaining the genome of HRTV-DL we established that the virus is present in Deep Lake by blasting the virus genome against available metagenomes from 2006, 2008, and 2013-15 [5] in the Integrated Microbial Genomes and Microbiomes (IMG) database [17]. *Hrr. lacusprofundi* was also detected in Deep Lake in the same metagenomes [5], confirming that both host and virus have been co-existing over several years.

### Genomic features of HRTV-DL1 and proteins associated with the virus particles

The genome of HRTV-DL1 encodes 49 putative open reading frames (ORFs) (Figure 2, Table S2). Using Hidden Markov models (HMM) [18] and domain prediction [19] we can predict the function for about 40 % of the annotated open reading frames (ORF) (Table S2). Two ORFs were identified that we propose to be involved in virus genome replication, both are also encoded by arTV CGphi46 (Figure 2). ORF48 is predicted as primase-helicase, probably generating a short primer that is then elongated by the host DNA polymerase, and ORF41, predicted as DNA polymerase sliding clamp, binding the DNA polymerase and preventing its dissociation from the template DNA. Two ORFs could be involved in transcription. ORF43, a predicted transcription initiation factor IIB, might be responsible for recruiting the host RNA Polymerase and ORF32, with weak hits to antitermination protein NusA, could associate with the host RNA polymerase elongation complex. ORF29 is a predicted adenine-specific methyltransferase and 100 % identical to a host protein, probably recruited by the virus to protect its genome from host R-M-systems. ORF7 and ORF8 were identified as small and large subunit of the terminase, responsible for genome packaging into the virus particle.

**Figure 2:**
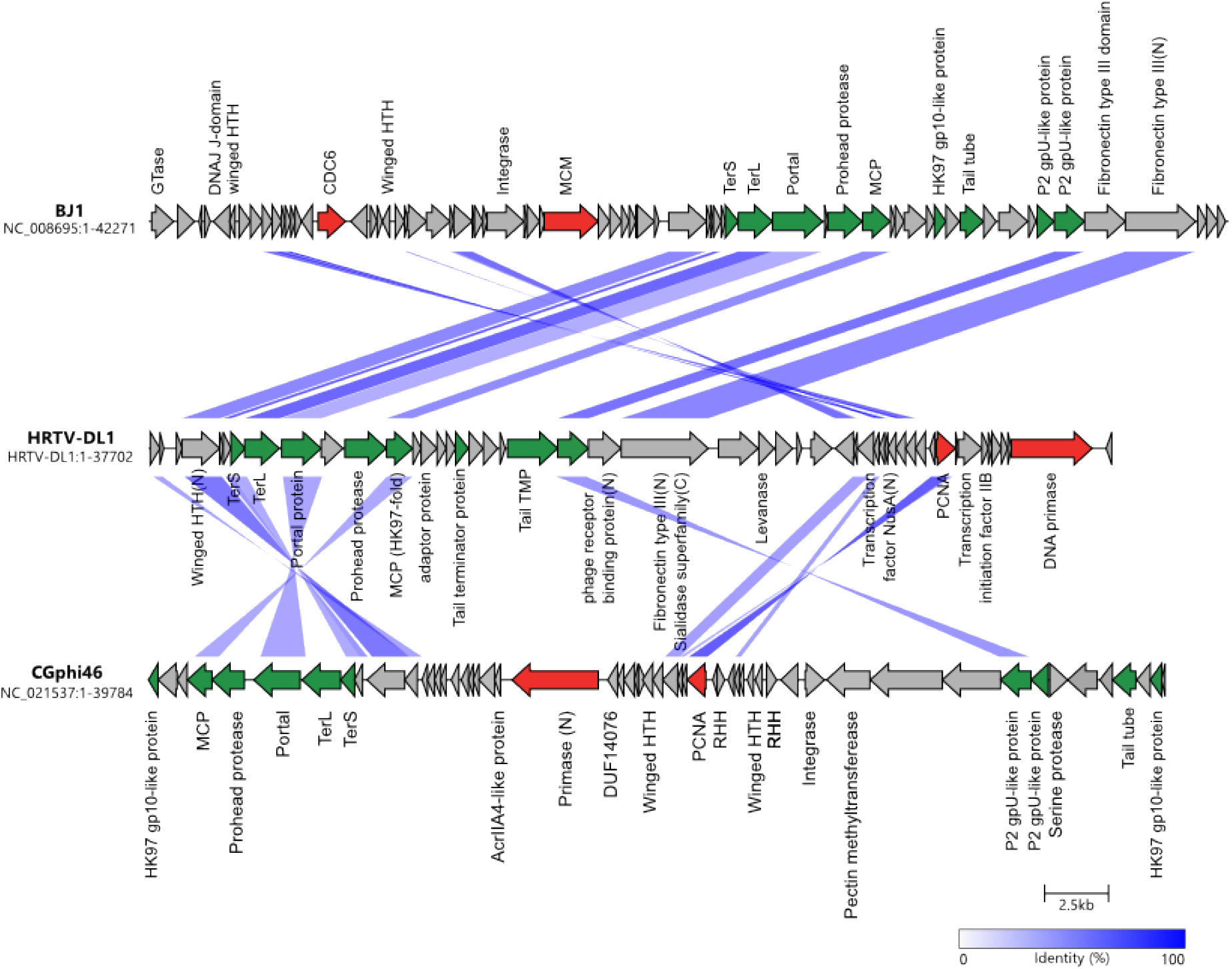
Comparisons of genomes HRTV-DL1 with BJ1 and CGphi46 belonging to the *Flexireviridae*. Protein functions are indicated above or below the corresponding ORFs. Genes encoding virus morphogenesis–related proteins are colored in green, whereas replication-related genes are colored in red. Homologous genes shared between viruses are connected by shadings of different degrees of blue based on the amino acid sequence identity. Figure was generated with clinker & clustermapjs [73].

Proteins building the virus particle were identified by mass spectrometry of purified virus particles and confirm our predictions (Table S2 and S3). The major capsid protein belongs to the HK97-like fold and is most similar to that of arTV BJ1 (45 % identity). Additionally, we could identify the phage portal protein, two capsid proteins with similarity to *Natrialba* phage PhiCh1 capsid proteins, the putative head-tail adaptor, the tail tape measure protein and the tail terminator protein. ORF18 and ORF25, identified in the virus particle, did not show any similarities to other viral proteins, therefore, their role within the virus particle remains enigmatic. Two proteins that could be involved in host attachment were identified. ORF24, that is present in virus particle, exhibits a immunglobulin-like fold (fibronectin type III domain) at the N-terminus. This domain is commonly found in bacterial phage tails and has been shown to exhibit membrane binding activity [20–22]. In the C-terminus we identified a sialidase domain. Sialidase (neuraminidase) is known to cleave glycosidic linkages and is essential for some eukaryotic viruses [23]. It could be involved in destabilizing the glycosylated S-layer and facilitating access to the host membrane. ORF25, also detected in virus preparations, shows similarities to a phage receptor binding protein with a high probability (P= 99.06). ORF4, with significant similarities to DNA double-strand break repair protein MRE11, and ORF43, annotated as transcription initiation factor, were also detected in virus particles. Both could be associated with the virus genome. ORF4 might facilitate circularization of the virus genome within the host, while ORF43 could recruit the host RNA polymerase ensuring timely transcription of the viral genome after injection.

Finally, we attempted to identify proteins involved in cell lysis. The mechanism of egress by complete membrane disruption, as it is known for head-tailed viruses of bacteria, has so far not been elucidated for arTV [24]. Lysis in gram negative bacteria is usually induced by two proteins. A holin that permeabilizes the lipid layer and an endolysin that subsequently degrades the cell wall [25]. Haloarchaea don not exhibit a peptidoglycan cell wall, but are protected by a protein shell, the S-layer. Specific proteases degrading the S-layer have not been found yet [24] and we could also not identify a protease on the HRTV-DL1 genome. However, glycosylation plays a significant role for the stability of the S-layer in haloarchaea [26] and degradation of the glycan can lead to a destabilisation. ORF26 shows similarity (using Hidden Markov models [18]) to a levanase and other glycosyl hydrolases with a high probability (P=99.12, 98.84 respectively) and could be responsible for glycan degradation and S-layer destabilization. Identifying holins is difficult due to their sequence diversity and only a few putative holins have been detected in archaea, but none on arTV genomes [27]. Holins are typically small proteins (75-174 aa) with 1-4 transmembrane domains (TM-domain). We can identify 3 small proteins with TM domains on the HRTV-DL1 genome, ORF6 (100 aa, two TM domains), ORF20 (67 aa, one TM domain) and ORF27 (222 aa, one Signal peptide, one TM domain). ORF6 and ORF27 hit other arTVs with low identity, however, none of the 3 ORFs show any similarities to proteins in public databases. Most holins are encoded by the gene adjacent to the endolysin gene in bacteria, making ORF27 to a potential candidate for a holin. Protein structure prediction of ORF27 [28] and comparison with the PDB database did not reveal any significants hits, however, only one holin structure has been determined so far [29]. Therefore, a potential involvement of ORF26 and ORF27 in cell lysis will need experimental investigation.

Genome comparison with 63 arTV (archaeal tailed viruses) [13] and phylogenetic tree construction (Figure S5), suggests HRTV-DL1 to be closest related to arTV BJ1 [30] and CGphi46 (NC_021537) (Figure 2) that have been classified into the *Flexireviridae* family. However, HRTV-DL1 is sufficiently different to be classified into a different genera, that we propose to name *Deelavirus* in accordance with its origin (**Dee**p **L**ake).

### Life cycle of HRTV-DL1

To gain insights into the initial stages of HRTV-DL1 entry, we followed the adsorption of HRTV-DL1 to ACAM34 host cells. The adsorption was very rapid, with ~75 % of virions being bound to cells within the first 30 seconds of infection. Further adsorption plateaus after 15 min with 99 % of particles bound to host cells (Figure S6). Such a rapid adsorption is rather uncommon for haloarchaeal arTV. Haloarcula hispanica tailed virus 1 (HHTV-1; *Siphoviridae*), binds extremely slowly with virions adsorbed only after 3h and binding of *Halorubrum* virus HRTV-1 was only detected after 25min [31]. Such diversity in adsorption rates indicates differences in adsorption mechanisms and receptors. The rapid adsorption of HRTV-DL1 suggests that the receptor is easily accessible and highly abundant on the cell surface. After successful infection, host cell lysis occurred between 56 and 68 hours post infection (p.i.) at 28 °C. The virus-to-host ratio (VHR) was determined by qPCR as 95 viral genome copies per host main chromosome copy number a few hours prior lysis (Figure 3). Interestingly, we observed changes to the cell morphology of infected *Hrr. lacusprofundi* at the onset of cell lysis. While the majority of the uninfected cells retained their rod-shaped morphology, infected cells tended to round up in the late stages of infection (Figure S7). However, a significant increase in cell size, as shown for another archaeal virus [32], was not observed.

**Figure 3:**
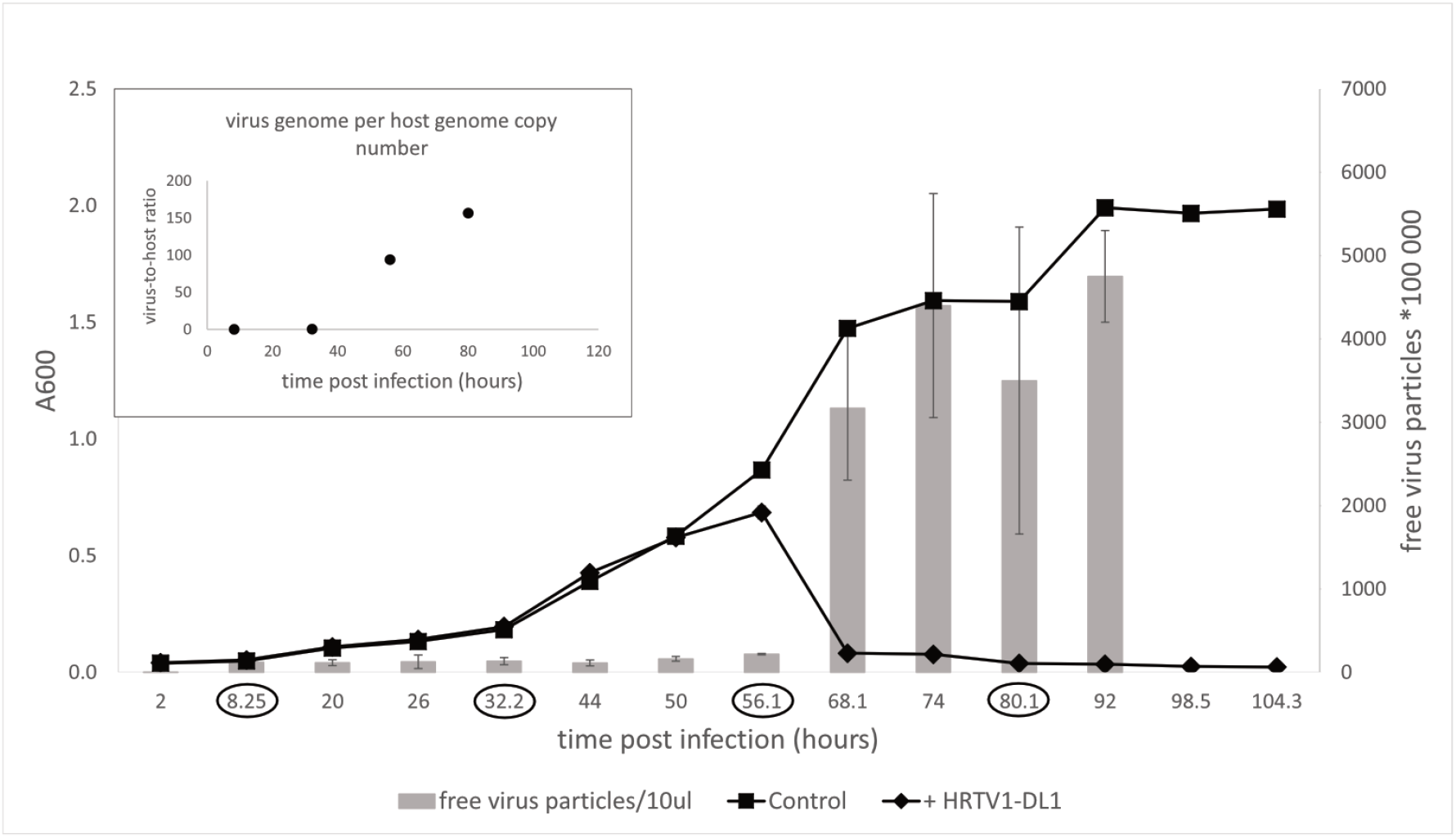
Life cycle of HRTV-DL1. Growth curve of uninfected control and HRTV-DL1 infected *Hrr. lacusprofundi* ACAM34_UNSW. Free virus particles in 10ul culture supernatant were determined by plaque assay. **Inlet:**Virus-host ratio determined by comparing copy numbers of host main chromosome and virus genome. Copy numbers were determined per ml cell culture by qPCR and samples were taken at time points encircled in the growth curve in the main figure. Graphs represent one of three biological replicates.

Both *Hrr. lacusprofundi* and HRTV-DL1 were isolated from a lake that experiences very low temperatures [1]. To determine whether low temperatures have an influence on the virus life cycle, we performed growth experiments at different temperature between 4 °C and 30 °C. Host growth rates change at different temperatures, consequently, lysis was observed at different time points post infection. However, lysis always occurred in early exponential phase at all temperatures (Figure S8). Therefore, we conclude that temperature does not influence the life cycle of HRTV-DL1 under the laboratory growth conditions tested. However, the presence of integrases in the HRTV-DL lysate indicate that other virus variants could exhibit a lysogenic life cycle, and this life cycle could well be influenced by temperature in its natural environment.

### HRTV-DL1 exhibits a very narrow host range that uncovers genetic differences in the same ACAM34 strain from different sources

HRTV-DL1 was isolated from Deep Lake, that is dominated by 4 different haloarchaea (*Halohasta litchfieldiae* tADL, *Hrr. lacusprofundi*, halophilic archaeon DL31, and *Halobacterium* DL1) [3]. CRISPR spacers against HRTV-DL1 were detected in the CRISPR loci of all except 1 species (*Halobacterium* DL1) (Table S4). The presence of spacers against HRTV-DL1 in the CRISPR loci of *Hrr. lacusprofundi* indicate that the virus, or variants of it, have been present in Deep Lake already before 1988. CRISPR spacers of all three different Deep Lake haloarchaea target different positions on the virus genome, indicating that they were acquired independently and not by gene transfer. Therefore, DL31 and tADL could be potential host organisms for HRTV-DL1. We tested all 4 species, one other *Hrr. lacusprofundi* strains from Deep Lake (DL18) and *Hrr. lacusprofundi* R1S1 from Rauer Lake 1, Antarctica [5, 8], but only *Hrr. lacusprofundi* ACAM34 was susceptible to infection. Adsorption assays revealed that attachment of the virus to all tested strains is impaired (Figure S9), showing that infection is already compromised at the adsorption stage. We assume that the attachment site of HRTV-DL1 is highly diverse, even in different strains of the same species. Modifications of cell surface proteins, in particular the S-layer that was found to be highly distinct, has already been proposed as a virus exclusion mechanisms of Deep Lake haloarchaea [5, 10].

When continuing characterisation of HRTV-DL1 in a new laboratory, we used a fresh stock of *Hrr. lacusprofundi* ACAM34 provided by the DSMZ. Surprisingly, while the virus successfully adsorbed to the host cell and injected its genome (Figure S9), HRTV-DL1 was not able to complete a lytic life cycle in the DSMZ strain. No lysis was observed in cultures and virus DNA could not be detected within the host at 56 hours p.i. (Figure S10). We subsequently assumed that the HRTV-DL1 susceptible strain, from now on referred to as ACAM34_UNSW (for University of New South Wales, Australia, the location of the laboratory the strain originated) had experienced genomic changes while being grown over a long time in the laboratory.

### Genome comparison of the ACAM34 laboratory strain with the ACAM34 strain from a culture collection reveals dramatic genomic rearrangements in the laboratory strain

To determine genetic differences between the DSMZ strain (ACAM34_DSMZ) and the HRTV-DL1 sensitive strain (ACAM34_UNSW), we re-sequenced the genome of both strains.

ACAM34_DSMZ assembled into 4 contigs. Contig 1 is circular with a size of 2,735,247 bp, (coverage 1080) representing the main chromosome (CHR1) with 99.99 % identity to the published genome. Contig 2 is circular with a size of 525,899 bp (coverage 1167), representing the secondary replicon (CHR2) with 99.99 % identity to the published sequence. Contig 3 is circular with a size of 431,344 bp (coverage 1137), representing the plasmid pHLAC01 with 99.99 % identity to the published sequence. Contig 4 is linear with the size of 6,176 bp matching a short region on CHR2 covering Hlac_3232 to Hlac_3234 with an insertion of a transposon within Hlac_3234. This contig has a lower coverage (432) than contig 2 and likely represents a variant of this region in the secondary replicon of the strain. In conclusion, the genome of ACAM34_DSMZ is to 99.99 % identical with the published sequence.

Surprisingly, ACAM34_UNSW assembled into only one single circular contig of 3,058,421 bp. Genome comparison with the published genome (Figure 4 and Figure 7) revealed that the entire plasmid pHLAC01 and about 38 % (173 kb) of CHR2 is not present, while the remaining 61.11 % of CHR2 (Hlac_2782 - Hlac_3106) is integrated into CHR1 between Hlac_1757 and Hlac_1759, two copies of ISH3 family transposase ISHla1 [33] with 100 % sequence identity. Hlac_2781 and Hlac_3106 flanking the CHR2 region integrated into CHR1 are also ISHla1 transposases that are 100 % identical to Hlac_1757 and Hlac_1759. We suggest that the integration of CHR2 occurred by homologues recombination between the copies of ISHla1 and at the same time caused the loss of the remaining CHR2. The insertion of a large genomic fragment into the main chromosome due to transposition activity has already been reported for a *Halobacterium salinarium* laboratory strain [34], and the insertion of a large plasmid into the main chromosome has been reported for *Haloferax volcanii* [7]. Apart from the insertion of CHR2, CHR1 remains highly conserved with an identity of 99.999 %. One minor change is found in an intergenic region between Hlac_0599 and Hlac_0600 that is missing a stretch of 23 nucleotides. The integrated remains of CHR2 have three significant changes. First, a transposase was inserted into Hlac_2793, a predicted ADP-ribosylglycohydrolase, and destroyed the gene. Second, Hlac_2835, with a MarR-like HTH domain within the N-terminus, has a number of changes on amino acid level within the C-terminus. And third, one of the two 16s ribosomal RNA genes has a number of nucleotide changes that make up 3% of the sequence. It remains unknown how the plasmid pHLAC01 was lost in ACAM34_UNSW. The secondary replicons are thought to be important for adaptation to varying environmental conditions. However, when grown under defined laboratory conditions the plasmid does not seem to be essential and is therefore likely too costly to be maintained. pHLAC01 exhibits the only CRISPR locus on the ACAM34 genome. This CRISPR locus contains 10 spacers that give a 100% match to the HRTV-DL1 genome. We therefore assume, that the resistance of ACAM34_DSMZ against HRTV_DL1 infection could be caused by the intact CRISPR system on pHLAC01, that is lost in ACAM34_UNSW. Nevertheless, there are also a number of other putative virus defense systems encoded on pHLAC01 and the region of CHR2 that is lost in ACAM34_UNSW, such as restriction modification systems (R-M-systems), Argonaut proteins and a BREX system (Table S5). These could also cause ACAM34_DSMZ resistance to HRTV-DL1 infection. Only two putative defense mechanism, one Argonaut protein (ACAM34UNSW_01791, Hlac_2785) [35], is still present in ACAM34_UNSW, leaving the strain basically unprotected against viral infection without any known virus exclusion mechanism.

**Figure 4:**
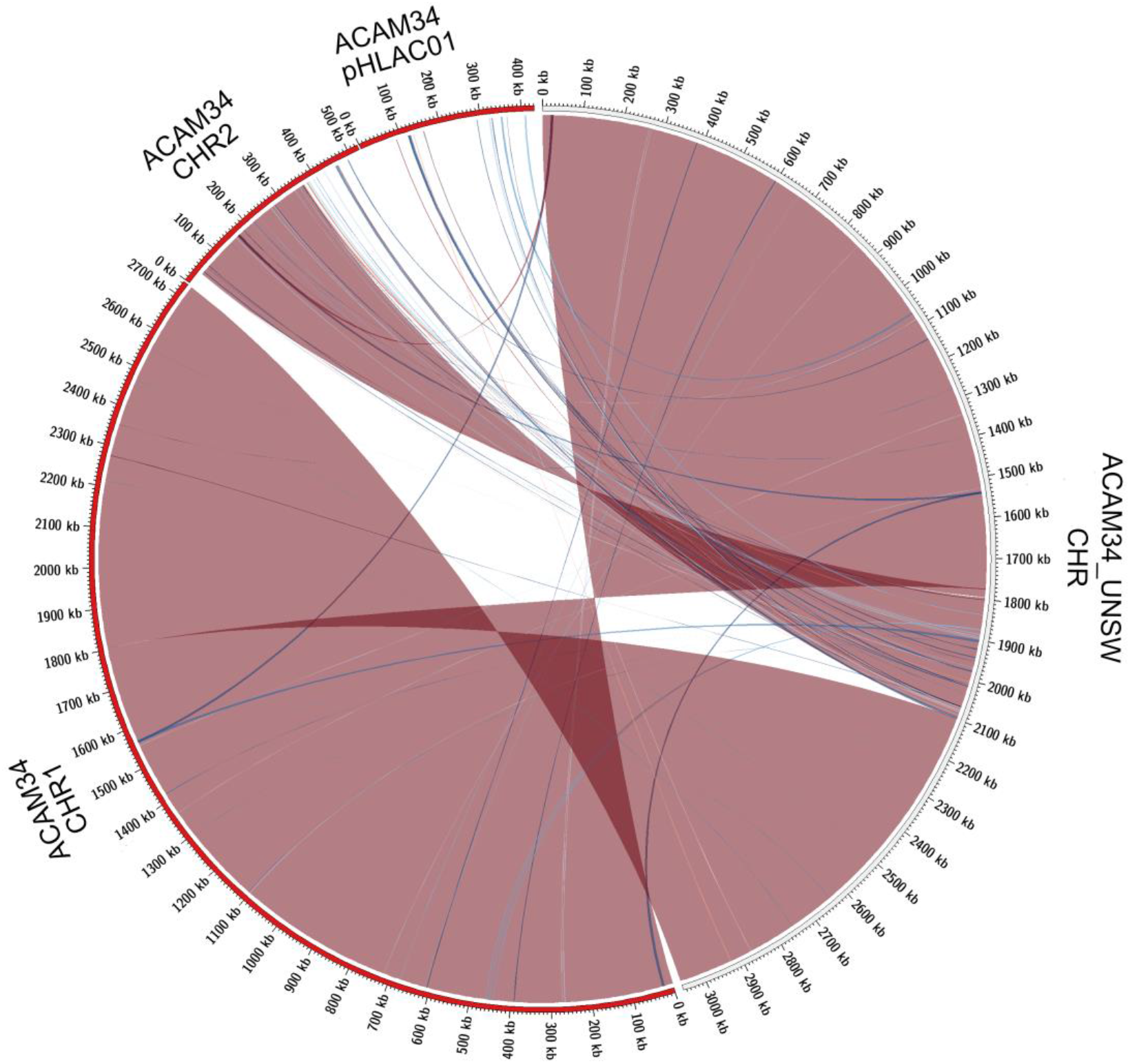
Genome comparison of ACAM34_UNSW with the three replicons CHR1, CHR2 and pHLAC01 (NC_012029.1, NC_012028.1 and NC_012030.1) of the reference genome from NCBI (ATCC 49239). Interconnecting lines highlight regions present in both genomes (red/blue colors indicates same/reverse orientation on the two genomes). The figure highlights that the ACAM34_UNSW genome consists of a single replicon comprising the full length primary replicon (NC_012029.1) and parts of one secondary replicon (NC_012028.1) of the reference genome, while pHLAC01 (NC_012030.1) of the reference genome is lost. Genomic regions shared between the two genomes were identified using NUCmer [74] and visualized with Circos [75] wrapped within the script mummer2circos.py (https://github.com/metagenlab/mummer2circos).

### Response of ACAM34_DSMZ to HRTV-DL1 infection does not show increased activity of any of the predicted virus exclusion mechanisms

To determine the mechanism underlying the resistance of ACAM34_DSMZ to HRTV-DL1 infection, we analysed transcriptional changes at two time points 32 hours post infection (middle phase, after infection is usually established in ACAM34_UNSW) and 56 hours post infection (late phase, immediately before ACAM34_UNSW is usually lysed).

The ACAM34_DSMZ transcriptome recovered no reads for the virus genome in both samples, which is consistent with our results that the virus genome copy number is already very low at 32 hours p.i. (time point 1 =T1) and not detectable at 56 hours p.i. (time point 2 =T2) (Figure S10). However, when comparing uninfected controls with infected samples, we still detect some changes to the host transcription. At T1 (32 hours p.i.) 468 genes are upregulated more than 2 fold (log2>1) and none are downregulated, and at T2 (56 hours p.i.) we detect 98 genes up- and 5 downregulated (Table S6). When analysing the functional potential of the upregulated genes at both time points, the majority falls into the categories transcription, replication and repair mechanisms, posttranslational modification, nucleotide metabolism and energy production and conversion (Figure 5). Interestingly, within the 30 most upregulated genes at T1, we identified 4 ORFs annotated as cold-shock proteins (CSPs) (Hlac_1630, Hlac_0048, Hlac_1986 and Hlac_1312). However, none of them is upregulated at T2, so this appears to be an early stress response to virus infection. In contrast, within the 30 most upregulated genes at T2 we identified three ORFs annotated as universal stress proteins (UspA) (Hlac_0044, Hlac_1730, Hlac_1573), possibly representing a late stress response to virus infection. The most upregulated gene at T1 is Hlac_2176 a predicted ferredoxin. Ferredoxins are small electron transfer proteins that are ubiquitous in biological redox systems [36] and upregulation of might be a response to oxidative stress caused by the virus infection. However, Hlac_2176 shows 87% sequence identity to a ferredoxin in *Hbt. salinarium* (OE_4217R), that was shown to be a coenzyme of alpha-ketoacid oxidoreductases [37], suggesting a possible involvement in energy conversion. Second most upregulated gene is Hlac_1324, that contains a winged helix DNA-binding domain and is likely a transcriptional regulator. Other genes within the 30 most upregulated genes are some ribosomal proteins and two other putative transcriptional regulators. While there is some basal expression, no significant transcriptional changes (log2>1) were observed for the integrated provirus Hlac-Pro1, suggesting that the integrated virus does not respond to superinfection, while such a response has been shown for proviruses in *H. volcanii* [12].

**Figure 5.**
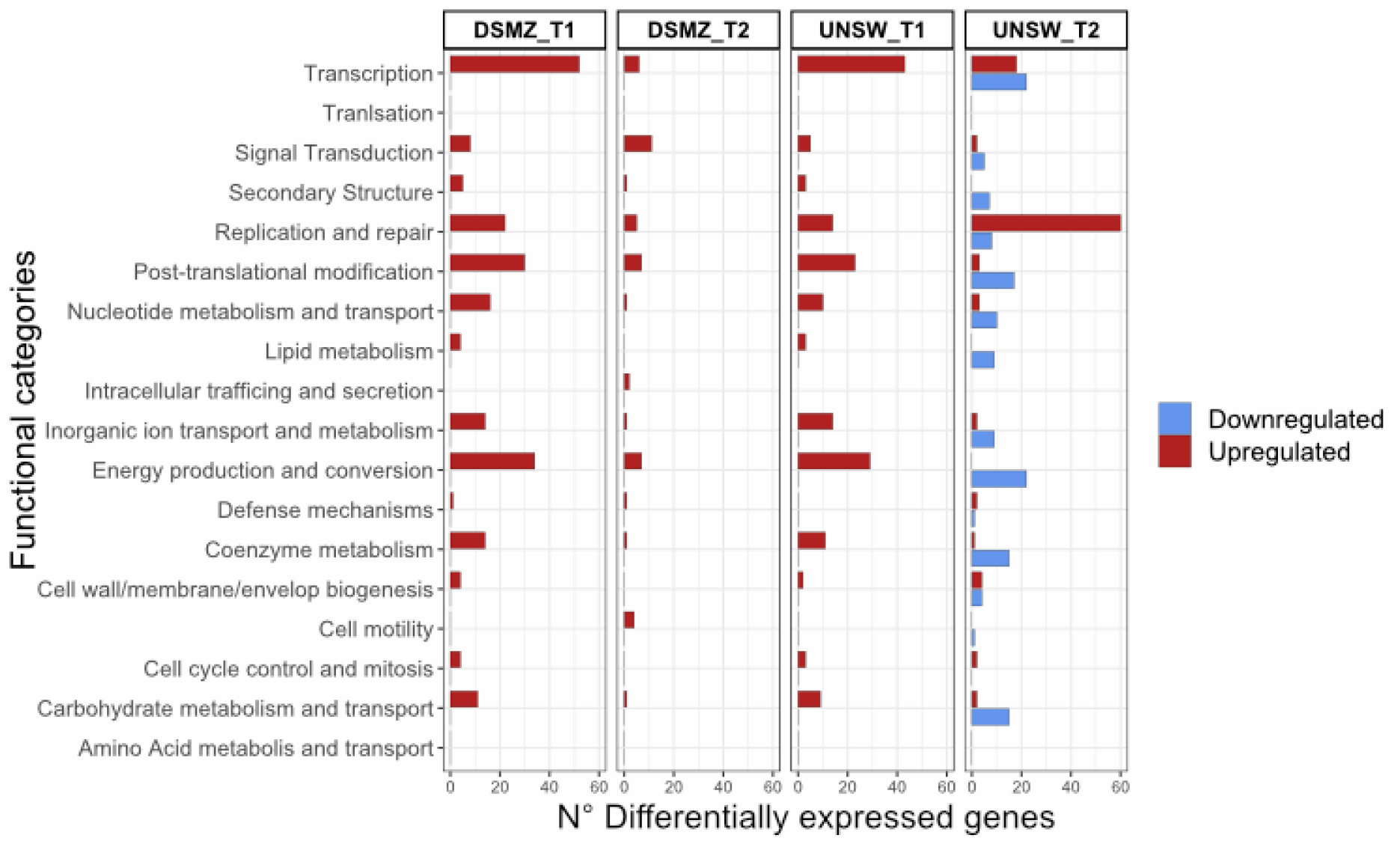
Functional profile of differentially expressed genes in ACAM34_DSMZ and ACAM34_UNSW under infection with HRTV-DL1. Bars represent the number of differentially expressed genes assigned to each particular functional category at a given time point (T1 = 32 hours post infection, T2 = 56 hours post infection). Functional classification of the *Hrr. lacusprofundi* genomes was performed using the cluster of orthologous genes (COG) database. The colors of the bars indicate if genes are upregulated (red) or downregulated (blue) respectively. Genes that were not assigned to any functional category are not displayed on the plot (for detailed information see Table S6 and S8).

Only one gene categorized as defence mechanisms is upregulated (Hlac_0998), however, the gene is not predicted to be involved in virus defence mechanisms (Na -driven multidrug efflux pump). Despite the presence of a number of spacers against HRTV-DL1 in ACAM34_DSMZ CRISPR loci (Table S4), none of the CRISPR-Cas genes showed upregulation (log2>1) at the two time points. Cas6b (Hlac_3333) is slightly upregulated with log2=0.5 at T1 and Cas6d (Hlac_3572) with log2=0.75 at T2, which could indicate some activity of the CRISPR-Cas system against HRTV-DL1. None of the other predicted virus defence mechanism (Table S5) show significant upregulation (log2>1). Perhaps, the basal activity of the CRISPR-Cas system, or basal activity of any of the other systems, is sufficient to exclude HRTV-DL1 (Figure 6). Alternatively, upregulation might have only been detectable shortly after virus genome injection (before 32 hour p.i.), even though other studies observed upregulation over an extended period during virus infection [38]. However, it is also possible that neither the CRISPR-Cas system, nor the other systems, are the main driver of HRTV-DL1 exclusion.

**Figure 6.**
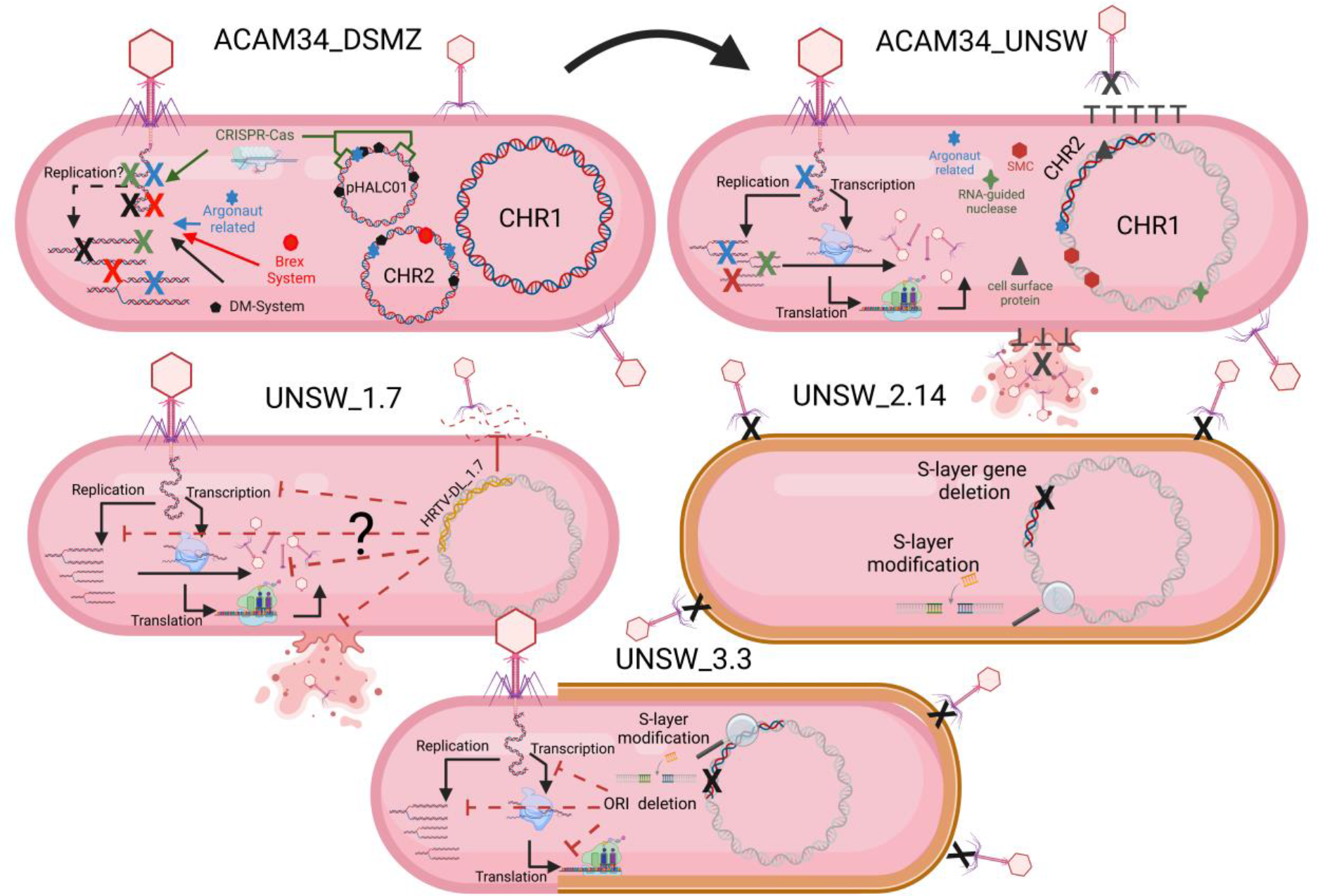
Summary of interactions observed between *Hrr. lacusprofundi* ACAM34 and HRTV-DL1. *Hrr. lacusprofundi* ACAM34_DSMZ allows adsorption and genome injection of HRTV-DL1. However, The genome is either degraded immediately after injection or replication is inhibited by one or several virus exclusion mechanisms present in the strain. ACAM34_DSMZ lost pHLAC01 and a portion of CHR2, while the remaining CHR2 is integrated into CHR1. All virus exclusion mechanisms potentially active against HRTV-DL1 in ACAM34_DSMZ are lost. However, other virus exclusion mechanism are activated allowing the emergence of escape mutants. Escape mutant UNSW_1.7 includes an integrated HRTV-DL1 variant, that provides superinfection exclusion. Escape mutant UNSW_2.14 lost one of two S-layer genes while the second S-layer gene experienced an insertion of two amino acids. This leads to a change of the cell surface and prevents HRTV-DL1 adsorption. Escape mutant UNSW_3.3 experienced an insertion of 12 amino acids into one of the two S-layer genes in 90% of the population, preventing adsorption of HRTV-DL1 to the majority of cells. The deletion of an origin of replication (ORI) and the subsequent repurposing of CDC6 is hypothesized to prevent virus particle production by a so far unknown mechanisms. Figure created with BioRender.com.

Therefore, we endeavoured to identify other potential HRTV-DL1 exclusion mechanisms. We also screened for upregulated genes that are not present on the genome of susceptible strain ACAM34_UNSW. Six genes were found to be upregulated on the portion of CHR2 that is absent in ACAM34_UNSW and 23 genes are upregulate on pHLAC01 (Table S7). The respective ORFs include two transposases, two predicted transcriptional regulators, a cdc6/orc1 replication initiation protein, a predicted universal stress protein, a few predicted RNA binding proteins and a number of small hypothetical proteins. Transposons are known to be activated upon virus infection [39] and co-localize with defense islands [40], they also have been found on virus genomes [13] and might inactivate viruses by insertion into essential virus genes. The universal stress protein and the transcriptional regulators are likely a host response to the stress condition rather than a virus exclusion mechanisms. A possible involvement of CDC6 in virus exclusion is discussed below (*Characterisation of virus escape mutants shed light on virus-host interactions and reveal additional virus exclusion mechanisms*). The remaining RNA binding proteins and small hypothetical proteins could represent so far undetected virus exclusion mechanisms, such as toxin-antitoxin (TA) systems or proteins influencing viral RNA stability. However, this would need to be investigated experimentally. In conclusion, our approach did not allow us to identify with certainty the virus exclusion mechanisms that are responsible for HRTV-DL1 exclusion in ACAM34_DSMZ.

### HRTV-DL1 infection induces the re-mobilization of CHR2 in ACAM34_UNSW and uncovers alternative virus exclusion mechanisms

We analysed the host response to HRTV-DL1 infection by determining transcriptional changes at two time points 32 hours p. i. (middle phase, after infection has been established, time point 1 = T1) and 56 hours p.i. (late phase, immediately before lysis, time point 2 = T2). Reads mapping the HRTV-DL1 genome represented 0.2% of all mapped reads 32 hours p.i. and 41% of all mapped reads at 56 hours p.i., consistent with the virus genome copy numbers increasing over time (Figure 3).

The transcriptional profile at time point 1 is very similar to that of T1 in ACAM34_DSMZ with 371 genes upregulated (log2>1) and none downregulated, and functional categories transcription, replication and repair mechanisms, posttranslational modification, nucleotide metabolism and energy production and conversion being the most upregulated (Figure 5). Additionally, we find the same genes among the 30 most upregulated genes, encoding for cold-shock proteins, ribosomal proteins, the ferredoxin Hlac_2176 and transcriptional regulators, being upregulated in ACAM34_UNSW and ACAM34_DSMZ, indicating that both strains exhibit a very similar response to early virus infection.

However, T2 shows a different profile, with 201 genes upregulated and 333 genes downregulated. The most upregulated functional category is ‘replication and repair’, consistent with a takeover of HRTV-DL1 and the degradation of the host genome (Figure S11). Genes involved in metabolic processes seem to be subjected to downregulation. Amongst the 30 most upregulated genes, only two were also detected as upregulated in ACAM34_DSMZ upon virus infection (Hlac_0148, Hlac_0919). Surprisingly, the majority of genes on former CHR2 (ACAM34UNSW_01788 to ACAM34UNSW_02100) was upregulated (Figure S12, Table S8). Therefore, we assumed that this region might have been re-mobilized during virus infection and that the upregulation is due to an increased copy number of the secondary replicon. Indeed when comparing copy numbers of CHR1 with former CHR2 by qPCR in the same samples, we detect a slightly higher copy number for CHR2 in the late stage of infected samples (1.3x the copy number of the main chromosome) (Figure S11). We conclude that CHR2 is mobilized only in a portion of cells in the population, because CHR2 usually has a copy number of 1.5-2 copies per CHR1 in ACAM34_DSMZ. When normalizing the expression values to the copy numbers, only 30 of 109 genes are still upregulated (log2>1), correcting the total number of upregulated genes from 201 to 122. As already observed in ACAM34_DSMZ, we do not detect differential expression of the integrated provirus.

Within the 30 most downregulated genes at T2 we find genes encoding for ribosomal proteins, for proteins involved in amino acid metabolism, proteins involved in energy production and conversion and proteins involved in oxidative stress response (Table S8), being a typical signature for a virus driven takeover of the host cell metabolism [41, 42].

Only one gene, predicted to be potentially involved in virus defence (ACAM34UNSW_01791), an Argonaut related nuclease (PADLOC [43]) that is still present in ACAM34_UNSW, is slightly upregulated (log2=0.9) in ACAM34_UNSW, but not in ACAM34_DSMZ (Hlac_2785) at any time point. We suggest that ACAM34UNSW_01791 is activated in ACAM34_UNSW due to the lack of other defence mechanisms, however, its activity does not seem to interrupt the lytic cycle of HRTV-DL1. Whether it has an impact on the efficiency of infection or virus production remains to be determined. To identify other potential virus exclusion mechanisms we analysed the 30 most upregulated genes. We propose that some of these genes might be implicated in virus exclusion of ACAM34_UNSW, that has lost all currently known virus defence systems (Figure 6). The predicted functions of the upregulated genes are summarized in Table S8 and some of particular interest, are discussed below.

Surprisingly, we find an entire genomic region being upregulated (ACAM34UNSW_00934 - _00942/ Hlac_0915 – Hlac_0923), with 8 of 9 genes within the 30 most upregulated genes. The region includes a predicted integrase (Hlac_0916), a couple of small hypothetical proteins and is enclosed by a tRNA gene and a transposase (Hlac_0923), indicating that it might have been part of a mobile genetic element that is activated upon virus infection. The region includes both partners of a two-component regulatory system, that is known to sense environmental changes and mediate the cellular response [44]. Whether the system is beneficial towards the host or high-jacked by the virus [45] remains to be elucidated.

ACAM34UNSW_01107 (Hlac_1086), that is originally annotated as transposase ORFB (conserved domain), also shows significant similarities (HMM) to Cas14 proteins that have been shown to exhibit RNA-guided nuclease activity [46]. However, since the ACAM34_UNSW does not contain any CRISPR arrays, we suggest that Hlac_1086 activity is more similar to the Transposon-associated TnpB, that is guided by an RNA derived from a sequence upstream of the TnpB [47]. Interestingly, we do find sequence similarities of the upstream region of Hlac_1086 with the PATE (see paragraph *HRTV-DL1 gene expression during infection reveals presence of a PATE with strong activity*) encoded on the virus and host genome. In absence of any other virus defence mechanism in ACAM34_UNSW, this gene might play an important role as a virus exclusion mechanism.

Two proteins (ACAM34UNSW_01618 and ACAM34UNSW_01620) within the 30 most upregulated genes, enclose a predicted chromosome maintenance protein (SMC), ACM34UNSW_01619 (log2=1.1). Additionally, another predicted SMC, ACM34UNSW_01796, is also upregulated (log2=1.1). SMC complexes were found to interfere with virus replication in humans [48, 49]. Additionally, SMC-like proteins were found to be associated with the recently discovered Wadjet system, that has been shown to exclude circular foreign DNA in bacteria [50, 51]. ACAM34UNSW_01618, ACAM34UNSW_01619 and ACAM34UNSW_01620 could potentially represent an archaeal derivative of this system.

Finally, another interesting candidate for a virus exclusion mechanism is ACAM34UNSW_02085 (Hlac_3088) (within the 50 most upregulated genes when normalized by 1.3). The gene is a passenger gene of a transposon with transposase ACAM34UNSW_02084 (Hlac_3078), and could be mobilized similarly to other virus exclusion mechanisms [40] or antibiotic resistance genes [52]. Hlac_3088 exhibits a PGF-TERM domain, an archaeal protein-sorting motif recognized by an archaeosortase [53], and is predicted to be a non-cytoplasmic protein with a TM domain. HMM search showed significant similarities with the S-layer protein of *H. volcanii*, however, Hlac_3088 is too small to be an S-layer protein. When comparing Hlac_3088 expression levels with those of the S-layer protein (Hlac_2976), Hlac_3088 has usually lower expression levels compared to the S-layer. However, at T2 it actually exceeds the expression levels of the S-layer gene (Table S9). We suggest that Hlac_3088 is located at the surface of the cell and could protect the cell by hiding the receptor and interfering with the attachment of the virus, similar to TraT, encoded by the F plasmid of *Escherichia coli*, that interacts with OmpA and blocks phage adsorption [54]. Alternatively, it could stabilize the cell and prevent or at least delay cell lysis.

While none of the proposed mechanisms are fully effective against HRTV-DL1, because the virus is completing its lytic cycle, they might prevent the lysis of the entire population and allow virus escape mutants to evolve.

### HRTV-DL1 gene expression during infection reveals presence of a PATE with strong activity

Transcriptomic data from HRTV-DL1 infected ACAM34_UNSW cultures revealed that all annotated HRTV_DL1 ORFs were expressed, while ORF3 was, by far, the most transcribed ORF (Table S10). When examining the reads mapping ORF3, we discovered that there is a peak that expands upstream of the annotated gene covering a region of approximately 300 bp and the majority of reads mapped antisense to the predicted coding sequence. Upon blasting the sequence against public databases, we discovered that it matched 99 % (1 mismatch, 1 gap) to a region within the hosts genome and a region that was previously described in plasmid pR1SE infecting the same host [8]. Further inspection revealed that this region represents a PATE (Palindrome-Associated Transposable Element), with a length of 489 bp that expands over the gap between ORF2 and ORF3. Only the C-terminal end of the PATE shows high expression, which includes the N-terminus of predicted ORF3, and expression occurs almost exclusively in reverse direction. We assume that ORF3 is a pseudogene and expression within this ORF is due to the PATE activity. In *Halobacterium salinarium* a PATE has been shown to be active as ncRNA [55] and we imagine a similar function for this PATE. However, it is encoded on both the host and the virus, so it is difficult to predict whether its activity is beneficial for the host or the virus, and whether the virus acquired it from the host or vice versa.

### ACAM34_UNSW can develop resistance to HRTV-DL1 infection

To observe adaptation to HRTV-DL1 infection, we established long-term infected cultures, in triplicates, of both ACAM34_UNSW and ACAM34_DSMZ with an MOI of only 1 to allow a recovery of infected cultures. ACAM34_DSMZ remained virus free over several dilution cycles and no new CRISPR spacers could be detected by PCR of the leader adjacent region of the CRISPR locus (Figure S13). For ACAM34_UNSW lysis was observed as expected between 45 and 70 hours post infection and the cultures were maintained to allow the growth of HRTV-DL1 resistant clones. The cultures slowly recovered over a time frame of app. 200 hours. After transfer of the recovered cells in fresh media, the cultures experienced another long lag phase of about 120 hours before establishing a normal growth (Figure S14). Cells from the recovered cultures were plated on agar plates to obtain single colonies and for each biological replicate we tested 24 colonies for infection with HRTV-DL1 by PCR. In 14 of the 72 tested colonies we could still detect the virus by PCR (Figure S15), indicating that the majority of the cells were able to exclude the virus.

Ten virus-free colonies were then chosen from each replicate to test re-infection with HRTV-DL1, and surprisingly, each biological replicate showed a different behaviour. Ten colonies chosen from replicate 1 could all be re-infected with HRTV-DL1, indicating that virus exclusion was only temporarily and likely due the activity of an existing virus exclusion mechanisms. All tested colonies were lysed identical to the original ACAM34_UNSW, except one colony ACAM34_UNSW_1.7 showing a reduced lysis (Figure S16). In contrast, none of the colonies that were isolated from replicate 2 could be re-infected with HRTV-DL1 and showed normal growth behaviour, apart from one ACAM34_UNSW_2.10 that grew faster than all the other tested colonies (Figure S16). Finally, colonies retrieved from replicate 3 showed a mixed behaviour. 9 out of 10 could be re-infected with the virus, however, for 8 out of the 9 the PCR signal for HRTV-DL1 was significantly weaker than in the control and no lysis could be observed in any of the cultures (Figure S16). We conclude that the majority of escape mutants retrieved from replicate 2 and 3, in contrast to replicate 1, had undergone some genetic changes to exclude HRTV-DL1.

Out of each replicate we chose one particularly interesting colony for further characterisation. ACAM34_UNSW_1.7 for showing reduced lysis, ACAM34_UNSW_2.14 as a representative for a fully resistant clone and ACAM34_UNSW_3.3 for showing a reduced infection without cell lysis.

### Characterisation of virus escape mutants shed light on virus-host interactions and reveal additional virus exclusion mechanisms

In-depth characterization of virus escape mutants (details presented in Supplementary Results and Discussion), including genome sequencing, revealed further insights into the interactions between HRTV-DL1 and ACAM34_UNSW. A graphic summary of the results is presented in Figure 6.

ACAM34_UNSW_1.7 is partially resistant to HRTV-DL1 infection, both HRTV-DL1 adsorption and virus particle production are significantly reduced (Figure S17 and S18). Genome sequencing uncovered that a variant of HRTV-DL1 (HRTV-DL_1.7), that is incapable of independent replication and virus particle production, is integrated into the UNSW_1.7 genome. We infer that HRTV-DL_1.7 is responsible for the partial resistance of UNSW_1.7 to HRTV-DL1 infection by a superinfection exclusion mechanisms as previously described for another arTV [16]. We propose that HRTV-DL_1.7 interferes with virus adsorption by causing changes to the cell surface of the host. However, how the integrated virus impacts virus particle production remains to be elucidated.

ACAM34_UNSW_2.14 is fully resistant to HRTV-DL1 infection (Figure S18). Infection is already impaired at the adsorption stage (Figure S17), indicating that the receptor for HRTV-DL1 might have undergone changes in this mutant. Genome sequencing revealed that UNSW_2.14 experienced a large genomic rearrangement of former CHR2 that was caused by transposase activity and led to the loss of about 200 kb. The deletion includes one of two S-layer genes (Hlac_2976) encoded by *Hrr. lacusprofundi*. While the strain is growing slightly slower than the parental strain (Figure S18), we did not observe any morphological changes to the cells (Figure S23) and conclude that the second S-layer gene is sufficient to maintain cell morphology and stability. However, we detected also an insertion of two amino acids into the second S-layer gene (Hlac_0412) and therefore conclude that the S-layer is the primary receptor for HRTV-DL1. Variability of S-layer genes of ACAM34 was previously detected by two ‘omics’-based studies of Deep Lake, and suggested to be driven by an arms race between viruses and host [5, 10].

Adsorption of HRTV-DL1 to escape mutant ACAM34_UNSW_3.3 is greatly reduced, but not fully impaired (Figure S17). While the virus genome can be detected in host cells, no cell lysis can be observed and no virus particles are detected in culture supernatants (Figure S18). We conclude that adsorption is partially impaired, but virus particle production and cell lysis is inhibited. The genome of UNSW_3.3 reveals an insertion of 12 amino acids into one of the S-layer genes (Hlac_2976) in 90% of the population, being the cause for the reduced adsorption and confirming the S-layer as primary receptor for HRTV-DL1. Only one other genomic change is detected in UNSW_3.3, the deletion of an origin of replication (ORI), that could possibly be responsible for preventing virus particle production and cell lysis. ORIs are binding sites for CDC6/ORC1 proteins and the deletion of an ORI could redirect CDC6 causing the same effect as the upregulation of CDC6 proteins that we detect in infected ACAM34_UNSW cells. Studies have shown that CDC6 can bind dsDNA without sequence specificity [56], that dysregulation of CDC6 expression can lead to inhibition of virus replication [57] and that CDC6 is also able to recruit the RNA polymerase I for rDNA transcription initiation [58]. Therefore, we suggest that CDC6 could affect HRTV-DL1 replication and transcription. Alternatively, it could be activated by the viral PCNA [59] and lead to ATP depletion.

## Conclusion

In this study we present an in-depth characterisation of the virus-host interactions between the archaeal head-tailed virus HRTV-DL1 and its host *Hrr. lacusprofundi* ACAM34. HRTV-DL1 was isolated from a sample taken in 2014 from the same environment (Deep Lake, Antarctica) as *Hrr. lacusprofundi* ACAM34, that was isolated in 1988. HRTV-DL1 was retrieved from a lysate (HRTV-DL) that was likely a mixture of different virus variants, and most importantly, HRTV-DL exhibited integrase genes that were almost entirely lost upon isolation of the pure variant HRTV-DL1. By using plaque assays we forced a selection for a variant with a lytic life cycle. We conclude that the virus has a high genome plasticity and our work demonstrates that working with an isolated model virus-host systems does not reflect the entire spectrum of the respective virus-host interactions as they are occurring in their natural environment.

Despite the fact that the time of isolation of virus and host is at least 26 years apart, the virus was able to generate a successful infection on ACAM34. However, the ability of HRTV-DL1 to infect the laboratory strain (ACAM34_UNSW), in contrast to its inability to complete a full lytic cycle in the same strain obtained from the DSMZ (ACAM34_DSMZ), uncovered that the laboratory strain had undergone a massive genomic rearrangement and lost an entire megaplasmid, demonstrating genome plasticity also for the host and demonstrating that the lost genomic information is not essential when grown under laboratory conditions. ACAM34_UNSW lost a number of virus defence mechanisms including a CRISPR-Cas system, upon growth under non-challenging laboratory conditions. This is a great example to demonstrate, that resequencing laboratory strains should be a good common practice.

We were not able to determine with certainty the mechanisms that is responsible for HRTV-DL1 exclusion from ACAM34_DSMZ. However, having both strains in hand, allowed us to uncover a number of new putative virus exclusion mechanism, that were only activated in ACAM34_UNSW in absence of other defence mechanisms. Analysis of the HRTV-DL1 infected transcriptome revealed the activity of a TnpB/Cas14-like nuclease and a transposon passenger gene that could potentially prevent the virus from attaching to the host cell. Additionally, we propose that a predicted SMC protein, in complex with at least two other proteins, might represent a derivative of the bacterial Wadjet system. Isolation of escape mutants, uncovered that one of two S-layer protein is the preferred virus receptor, indicating that exhibiting two S-layer proteins could be a strategy of virus exclusion. HRTV-DL1 can provide partial superinfection exclusion when integrated into the host genome. Additionally, we propose that the origin binding protein CDC6 can be involved in virus exclusion by a so far unknown mechanisms. We observed that both the host and the virus actively undergo genomic rearrangements when interacting with each other, allowing both host an virus to adapt to each other.

Our study uncovers numerous virus exclusion mechanisms within one single organisms, that have likely been acquired over a very long time by a continuously ongoing arms race between viruses and *Hrr. lacusprofundi*. The establishment and characterization of this virus-host system will allow to observe this ongoing race both in the laboratory and in the environment for which metagenomics data are available. The new virus exclusion mechanisms discovered in this study can be investigated in detail and possibly be identified in other organisms. ACAM34_UNSW itself can be used as model to study virus exclusion mechanisms. It is susceptible to a variety of viruses, including HRTV-DL1 and others [8, 12, 60–62], and virus exclusion mechanisms can be re-introduced [63] to test their activity against diverse viruses.

## Materials and Methods

### Isolation of HRTV-DL, strains and culture conditions

HRTV-DL, formerly described as DLHTHV [5], was isolated from the supernatant of a colony that lysed upon propagation in liquid culture. Briefly, a sample, taken from Deep Lake (Antarctica) [1] in 2014, was used to generate enrichment cultures in liquid medium (culture medium and conditions see below). Enrichment cultures were plated on solid media and a single colonies picked into liquid media. The colony was identified as *Halorubrum lacusprofundi* by 16s rDNA sequencing using universal primers (21F and 1510R [64]). After lysis was observed in the culture, cells and cell debris were removed by centrifugation (4,500 x *g*, 30 min), the supernatant of the culture (lysate) was filtered through 0.2 um filters and used for infection of *Halorubrum lacusprofundi* ACAM34. *Halorubrum lacusprofundi* ACAM34 (DSM 5036), hereafter named ACAM34 _DSMZ, was obtained from the German Collection of Microorganisms and Cell Cultures (DSMZ). The parental strain of *Halorubrum lacusprofundi* ACAM34_UNSW was provided by either Peter Franzmann or John Bowman (not traceable anymore) and subsequently cultured in the laboratory (Ricardo Cavicchioli, School of Biotechnology and Biomolecular Sciences, The University of New South Wales, Sydney) from glycerol stocks stored at −80 °C. The strain went through several rounds of culturing, plating and −80 °C storage in three different laboratories (UNSW and University of Technology, Sydney, Australia; Max Planck Institute for Marine Microbiology, Bremen, Germany) prior to re-sequencing. The number of generations that the strain had undergone during this time is not traceable. *Halohasta litchfieldiae* tADL (DSM 22187) was provided by the DSMZ. Halophilic archaeon DL31 and *Halobacterium* DL, and *Hrr. lacusprofundi* R1S1 and DL18 were isolated previously [3, 8]. All strains were grown in DBCM2 media [61], with 5 g peptone and 1 g yeast extract added per liter. Incubation temperature was 28 °C unless stated otherwise. Cultures were incubated in glass flasks aerobically at 120 rpm. Solid ager plates contained 16 g agar per liter and top layer agar contained 6 g agar per liter. Plaque assays were performed by mixing virus and host with 10 ml of top layer agar that was subsequently poured on solid plates and incubated at 28 °C until growth was visible.

### Isolation and purification of HRTV-DL1 particles

For virus production, *Hrr. lacusprofundi* was grown in liquid culture to mid exponential phase, with an optical density at 600 nm (OD600) of 0.5. Cells were harvested (4,500 x *g* for 45 min), mixed with viral suspension with a multiplicity of infection (MOI) of 5-10 and incubated for 2 hours at room temperature to allow viral adsorption. Treated cells were inoculated into 500 ml liquid cultures that were monitored by measuring optical density changes (OD600). After lysis occurred, cultures were centrifuged at 4,500 x *g* for 45 minutes to pellet the cells. The supernatant was recovered and viruses were subsequently precipitated with polyethylene glycol (PEG) 6,000 (10 % w/v final concentration) and incubation at 4 °C overnight. Then, viral preparations were collected by centrifugation (30,000 x *g*, 45 min, 4 °C). Pellets were resuspended in DBCM2 salt solution (DBCM2 media without nutrient sources pyruvate, trypton and yeast extract), sterile filtered (pore size 0.2 μm) one time when used for plaque assay and twice when used in liquid culture infection assays. Virus solutions were stored at 4 °C and were active for a minimum of 1 year. For downstream analyses such as genome sequencing and mass spectrometry the virus particles were further purified by Cesium chloride (CsCl) density gradient centrifugation. The virus solution was treated with 200 U of DNase I and 50 ug/ml of RNase A to reduce host genomic contamination prior to loading on a CsCl density gradient (0.45g CsCl /ml in DBCM2 salt solution) and centrifuged at 38,000 rpm for 22 hours, 4 °C (SW41 Ti Swinging-Bucket rotor, Beckman & Coulter). Bands containing virus particles were extracted with a syringe, diluted in three volumes of DBCM2 salt solution and re-precipitated with PEG (final concentration 10 %, 4 °C, overnight). After centrifugation at 30,000 x *g* for 30 min the resulting pellet was washed twice with DBCM2 salt solution and stored at −20 °C before further processing.

### Imaging

For Transmission electron microscopy (TEM), virus containing solution was adsorbed for 5 min to carbon coated copper grids and stained for 1 min with 2 % uranyl acetate (w/v in water). Electron micrographs were generated using JEM2100 Plus at 200 kV acceleration voltage. For light microcopy, cells were fixed with 1 % glutaraldehyde for 1 hour at room temperature and then stored at 4 °C until imaging with a Zeiss AxioPhot microscope with AxioCam MRm.

### DNA extraction, manipulation and genome sequencing

Genomic DNA of all samples, if not otherwise stated, was extracted using genomic DNA extraction kit (Bioline, London, UK) according to the manufacturer’s instructions. Plasmid extraction was performed with ISOLATE II Plasmid Mini Kit (Bioline, London, UK) according to the manufacturer’s instructions. Nuclease digestions using specified restriction enzymes (New England Biolabs [NEB], 20 units) or exonuclease III (NEB, 5 units) were performed on 2 – 3 μg of DNA extracted from virus particles for 1 h at 37 °C. PCR reactions were performed with the Q5^®^ High-Fidelity DNA Polymerase (NEB) and contained 0.02 units/μL of DNA polymerase, primer concentration of 0.1 μM for both forward and reverse, 1X of Q5 Reaction Buffer and 1X Q5 High GC Enhancer. The following program was used: 5 min at 95 °C, followed by 35 cycles of 30 s at 95 °C, 30 s at 68 °C for annealing and 30 s at 72 °C for elongation. Digested DNA and PCR products were separated on 1 % agarose gels in Tris-borate-EDTA buffer and stained with SYBR™ Safe DNA stain (Invitrogen). For HRTV-DL, library preparation (Nextera XT) and sequencing (MiSeq) was performed at the Ramaciotti Centre for Genomics (UNSW, Sydney, Australia). For HRTV-DL1, UNSW_1.7, UNSW_2.14 and UNSW_3.3 library preparation (FS DNA Library, NEBNext^®^ Ultra™) and sequencing (Illumina HiSeq3000, 2 x 150 bp, 1 Gigabase per sample) was performed at the Max Planck-Genome-Centre Cologne (Cologne, Germany). For PacBio sequencing of ACAM34_UNSW and ACAM34_DSMZ, DNA extraction, library preparation, sequencing (Pacific Biosciences Sequel, 4 samples on one SMRT cell) and assembly (‘Microbial Assembly’ function in PacBio SMRT^®^ Link v8.0 for ACAM34_DSMZ and flye assembly tool v2.6 [65] for ACAM34_UNSW) was performed at the Max Planck-Genome-Centre Cologne (Cologne, Germany).

### Genome assembly, annotation and phylogenetic analysis

After assembly with SPAdes [66], the HRTV-DL genome was manually closed to one contig using primers listed in Table S1, followed by Sanger sequencing (MICROMON DNA Sequencing Facility, Monash University, Victoria, Australia) of PCR products. The HRTV-DL1 genome was assembled using metaviralSPAdes [67]. Genome annotation for all genomes was done using Prokka [68] followed by manual corrections using conserved domain based searches [19] or hidden Markov model (HMM) based searches [18]. For genome comparison HRTV-DL and HRTV-DL1 nucleotide sequences were compared using BlastN (standard settings) and comparison was visualized using Artemis comparison tool [69]. ORFs of HRTV-DL and HRTV-DL1 were compared using BlastP (standard settings) (Table S1). Phylogenetic tree reconstructions from protein sequences was done with VICTOR under optimal settings (formula VICTOR d6), as implemented at the DSMZ webserver https://victor.dsmz.de. Prediction of termini was done using PhageTerm [70].

### Protein content analysis

Protein content analysis of HRTV-DL1 particles was performed by mass spectrometry on purified virus pellets as described previously for plasmid vesicles and membrane vesicles [8]. Peak lists were generated using Mascot Daemon/Mascot Distiller (Matrix Science, London, England) and initially submitted to the database search program Mascot (Matrix Science). Search parameters were: Precursor tolerance 4 ppm and product ion tolerances ± 0.05 Da; Met(O) carboxyamidomethyl-Cys specified as variable modification; enzyme specificity was trypsin; 1 missed cleavage was possible; customized databases searched: *Hrr. lacusprofundi* ACAM34 and HRTV-DL. After genome sequencing of ACAM34_UNSW, results presented in Table S3 were obtained using the SEQUEST search algorithm with Thermo Proteome Discoverer™ 2.5.0.400 (Thermo Fisher Scientific) with the same settings searching customized databases *Hrr. lacusprofundi* ACAM34_UNSW and HRTV-DL1.

### Virus infectivity and kinetics

To study the life cycle, cultures of *Hrr. lacusprofundi* (all strains) were synchronized using an adaptation of the “Stationary phase method” [71]. The strain was scratched from a −80 °C glycerol stock into liquid culture and grown up to an OD600 ~ 1. Then a 20-fold dilution step in fresh media was performed (final OD600 = 0.05) and cultures were then regrown up to OD600 ~ 1. Iterative dilution and growth of the culture were repeated twice before considering a culture synchronized. For infection with HRTV-DL1, cells from synchronized cultures (OD600 ~ 1) were collected by centrifugation, resuspended in 1 ml of fresh media and infected with HRTV-DL1 virus with a MOI of 5. After incubation (2 h, room temperature), cells were transferred into liquid cultures and growth was monitored by optical density (OD600). Long-term cultures were established by continuously diluting infected cultures at OD600 ~ 1 to OD600 ~ 0.05. Infection was confirmed by PCR using primers HRTV-DLF and HRTV-DLR (Table S11).Viral titer in culture supernatants was quantified by plaque assay after removal of cells by centrifugation (11,000 x *g*, 10 min, room temperature [RT]). Intracellular virus titers and host chromosome copy numbers were quantified by qPCR. Briefly, samples of 2 ml culture in biological replicates were collected and pelleted (11,000 x *g*, 10 min, RT). Cell pellets were washed two times with 1 ml of fresh media and stored at −20 °C upon DNA or RNA extraction. Quantification of *Hrr. lacusprofundi* and HRTV-DL1 genome copy number were carried out using a CFX96 Touch Real-Time PCR (Bio-Rad Laboratories, Inc., Hercules, CA, USA) and the software CFX Manager™ Software. Primers are listed in Table S11. Each reaction (10 μL) contained 1X SsoAdvanced Universal SYBR™ Green Supermix (Bio-Rad) and primer concentrations as stated in Table S11. The following amplification thermal cycling program was used for both primer sets: 5 min at 95 °C, followed by 40 cycles of 30 s at 95 °C and 30 s at annealing temperature stated in Table S11, with readings taken between each cycle. Efficiencies of the assays were 95–102 %, with R^2^ values >0.99 for all assays. The specificity of the qPCR was confirmed by unique signals in melting curves and gel electrophoresis of PCR products.

### Adsorption assays and host range assessment of HRTV-DL1

For adsorption assays, 5 ml of cells at OD600 = 0.5 were harvested by centrifugation (4,500 × *g*, 30 min, RT) and resuspended in 1ml fresh medium. The cells were subsequently infected using a MOI of 5. At defined time intervals the adsorption was stopped by immediate centrifugation (10,000 × *g*, 5 min, RT).

The number of remaining free virus particles in the supernatant was determined by plaque assay. A cell-free control (only media) served as control. Three different strains of haloarchaea (*Halohasta litchfieldiae* tADL, halophilic archaeon DL31, and *Halobacterium* DL1 [3]) and two additional strains of *Hrr. lacusprofundi* (R1S1 and DL18 [8]) were tested to determine the host range of HRTV-DL1. CRISPR matches to HRTV-DL1 were identified by searching for CRISPR loci using CRISPRs finder [72] and blasting (BlastN) identified spacer against the HRTV-DL1 genome. Adsorption of HRTV-DL1 to the different strains was determined by adsorption assay as described above and the absence of HRTV-DL1 DNA within cells was confirmed by PCR on cell pellets 1 day and 5 days post infection.

### Transcriptomic analyses

RNA extraction of frozen cell pellets with 3 biological replicates for the infected and 2 biological replicates for the controls, was performed with the Direct-zol™ RNA miniprep Kit (R2051, Zymo Research). RNA concentration and integrity were assessed using Nanodrop DS-11 Spectrophotometer (DeNovix) according to the manufacturer’s instructions. Library preparation and sequencing was done at the Max Planck-Genome-Centre Cologne (Cologne, Germany). Briefly, ribosomal RNA were depleted prior to sequencing using the rRNA depletion Kit riboPOOL^™^, for *Haloferax volcanii* (88.36% identity to 16s rDNA sequences of *Hrr. lacusprofundi*), siTOOLs Biotech^®^. Libraries were prepared with library kit NEBNext^®^ Ultra™ II RNA Library Prep Kit for Illumina and sequencing was performed on an Illumina HiSeq3000 sequencer, following a 1 x 150 run. Read trimming and mapping was performed with the ‘Map to reference’ function (Mapper ‘Geneious RNA’) with medium-low sensitivity within Geneious Prime^®^ 2022.2.1. Expression values (FPKM values) were calculated using standard settings and comparison of expression levels were performed using DeSeq2 within Geneious Prime^®^ 2022.2.1 using default settings. Genes with p-values < 0.01 and a fold change of at least two times (log2FC ≥ 1 or ≤ −1) were considered to be differentially expressed (DE).

### Analysis of ACAM34_UNSW escape mutants

For the isolation of ACAM34_UNSW HRTV-DL1 escape mutants we established long-term infected cultures as described above (Virus infectivity and kinetics), in triplicates. Cells were only infected with an MOI of ~1 to allow a recovery of infected cultures. After lysis of ACAM34_UNSW the cultures were maintained to allow the growth of HRTV-DL1 resistant clones. Cells from the recovered cultures were plated on agar plates to obtain single colonies. HRTV-DL1 infection was determined by PCR as described above. HRTV-DL1 life cycle in escape mutants and adsorption of HRTV-DL1 to escape mutants was determined as described above. Isolation of genomic DNA and genome sequencing is described above. For genomic analysis of escape mutants, reads were mapped against ACAM34_UNSW and HRTV-DL1 using ‘geneious mapper’ with medium-low sensitivity and default settings and variants were called using the Geneious function ‘Find Variations/SNPs’ with default settings in Geneious Prime^®^ 2022.2.1. Variants with a coverage below 150 and a variant frequency below 75% were excluded from the analysis. HRTV-DL1_1.7 was assembled using MetaviralSPades [67].

## Data availability

The genomes of HRTV-DL, HRTV-DL1 and HRTV-DL1_1.7 are available on NCBI under the with accession numbers OP630574, OP630575, OP630576. RNA sequencing raw data were submitted to ENA-EMBL under project number: PRJEB56294. Raw data from genome sequencing of UNSW_1.7, UNSW_2.14 and UNSW_3.3 are available in the Sequence Read Archive under BioProject ID: PRJNA887929. The genome of ACAM34_UNSW is associated with the manuscript as supplementary data file.

## Acknowledgments

We thank the Max Planck-Genome-Centre Cologne (Cologne, Germany) for assistance with DNA and RNA sequencing. We thank Ingrid Kunze (MPI for Marine Microbiology, Bremen, Germany) for assistance with some of the experiments. We thank Tomas Alarcón-Schumacher and Bernhard Tschitschko for help with some of the figures. We thank the entire Archaeal Virology group at the Max-Planck-Institute for Marine Microbiology and Friedhelm Pfeiffer for critically reviewing the manuscript. We also acknowledge Friedhelm Pfeiffer and Mike Dyall-Smith for identifying the PATE and helpful discussions. Part of this work was supported by the Australian Research Council [DP170102576]. Mass spectrometry results were obtained at the Bioanalytical Mass Spectrometry Facility (BMSF) within the Analytical Centre of the University of New South Wales. Subsidized access to the BMSF is gratefully acknowledged. Finally, we want to thank the Max-Planck-Institute for Marine Microbiology and the Max-Planck-Society for continuous support.

## Authors Contributions

C.M. and D.T. performed a majority of the experimental work. S.E. performed some of the experiments, conceived and led the study, and performed the primary writing of the manuscript. R.C. led the initial phase of the study. L.Z and M.J.R performed mass spectrometry and analyzed data. All authors participated in the analysis and interpretation of the data and contributed to the writing of the manuscript.

