## Supplementary material for "In depth characterization of an archaeal virus-host system reveals numerous virus exclusion mechanisms": Supporting_Information.pdf

### Supplementary Results and Discussion

#### *Escape mutant ACAM34_UNSW_1.7 includes a variant of HRTV-DL1 and exhibits superinfection exclusion*

ACAM34_UNSW_1.7 was shown to be re-infected by HRTV-DL1, but did show reduced lysis. To determine the mechanism behind the reduced lysis of this strain, we characterised the relationship of HRTV-DL1 with UNSW_1.7 and analysed the genome. Adsorption assays showed that HRTV-DL1 can bind to UNSW_1.7, but adsorption was significantly less efficient than for the parental strain (Figure S17). As observed in the previous experiment, infection of UNSW_1.7 resulted only in a partial lysis of the culture about 50 hours post infection, with a clear difference to the parental strain that showed a total lysis (Figure S18). This difference was confirmed when analysing the amount of free virus particles in the supernatant of the lysed culture. While the parental strain showed a strong increase in free virus particles after lysis the number increased only slightly for UNSW_1.7, having only 8.3% of the free virus particles detected for the parental strain (Figure S18). Similar to the parental strain, we observed morphological changes of the infected cells at the onset of lysis by light microscopy. However, while we could only detect cell lysates for the parental strain in the late stages of infection, the infected culture of UNSW_1.7 retained a significant number of intact cells. Surprisingly, the surviving intact cells show a similar morphology as the infected wild type cells shortly prior cell lysis (Figure S19).

Subsequently, the genome of UNSW_1.7 was re-sequenced and read mapping against the host and the virus genome revealed a significant amount of reads for the virus genome, however, not the entire genome of HRTV-DL1 was covered (Figure S20a). A new assembly (metaviralSPAdes) recovered a circular fragment of 28.6kb with a similar mean coverage (248.761) as the host genome (268.078). While the order of the different coding regions has changed in this new virus genome, we also find five specific regions that are lost on the HRTV-DL1_1.7 variant genome (Figure S20b). The first region, covering ORF2-ORF8, includes the region that we used to target the HRTV-DL1 genome by PCR, which elucidates why we did not detect the virus initially in uninfected UNSW_1.7. Subsequently, we were able to detect the HRTV-DL1_1.7 in uninfected UNSW_1.7 by PCR with a different primer set (Figure S21a). While the majority of ORFs in this region are hypotheticals, the lack of ORF7, the predicted small subunit of the terminase, could have a major impact on virion production. The second region covers the C-terminus of ORF17 and the entire ORF18, both proteins detected in virus particles, that are possibly crucial for virion assembly. The next cluster of genes that we could not detect in HRTV-DL1_1.7 are ORF22-ORF24, of which ORF23 and ORF24 are predicted to be involved in host attachment. ORF28, hypothetical, and ORF29 the methyltransferase are missing. The third region is a major gap in ORF41, the predicted DNA polymerase sliding clamp, that could have an influence on genome replication. Finally, HRTV-DL_1.7 also lost the PATE. This loss might have been essential to allow integration into the host genome considering that Hlac_1086 (Cas14/TnpB) might be indeed active against HRTV-DL1. In summary, HRTV-DL1_1.7 is likely not capable of virion assembly and genome replication, concluding that it is a defective virus. Surprisingly, the reassembled HRTV-DL1_1.7 contains six additional ORFs, Table S12) that we neither find on the final HRTV-DL1 genome (Figure S20b), nor recruited reads from both the DNA and the RNA sequencing data (Figure S20c). However, when mapping reads from the HRTV-DL sequencing we detect a low coverage for these regions (Figure S20d), indicating that they derive from the original HRTV-DL and have been retained as a very minor fraction in the HRTV-DL1 population. One of the six ORFs encoded in these regions,

ORF_00033, is a predicted site-specific integrase and could be responsible for the integration of this particular virus variant.

We tested UNSW_1.7 for presence of the virus variant as an isolated genome. The genome could neither be isolated from the supernatant of UNSW_1.7 cultures, nor as circular genome from cells of UNSW_1.7, even under UV induction (Figure S21b). Considering the similar coverage between the host UNSW_1.7 and the virus within the strain HRTV-DL1_1.7, we conclude that the virus variant is integrated into the host chromosome.

Analysis of the UNSW_1.7 host genome revealed a few minor changes (Table S13). We detected a number of single nucleotide polymorphisms (SNPs) in intergenic regions (7 in total), one in the 23s rRNA coding region and two SNPs within coding regions without effect on the amino acid sequence. Three SNPs were detected within coding regions and lead to an amino acid substitution, however, they are also present in the other two escape mutants and we assume that they derive from the parental strain. Two changes within coding sequences lead to a frame shift. In ACAM34UNSW_01960, a hypothetical protein, a frameshift occurs within the middle of the protein and another frame shift was detected in the N-terminus of ACAM34UNSW_00139 (78% of the population), a predicted zinc/iron permease. Finally, a region of about 7000 bp (ACAM34UNSW_02044 - ACAM34UNSW_02050), did not recruit any reads. However, this region was not detected in any of the three analysed escape mutants, we confirmed by PCR that it is also not present in the parental strain and assume that it is an assembly error.

The isolation of a resistant escape mutant that incorporated the virus into its genome has already been described for *Halobacterium salinarium* virus ØH [1]. The study also describes genomic rearrangements of the virus genome upon incorporation, similar to what we have observed for HRTV-DL_1.7. The integration of this low abundant HRTV-DL1 variant also confirms that the exclusive lytic life style that we observe in the laboratory is not the only life style exhibited by the virus. Variants of HRTV-DL could be found integrated into host genomes in its natural environment. We suggest that the integrated HRTV-DL_1.7 is responsible for the partial resistance of UNSW_1.7 against re-infection with HRTV-DL1 by a superinfection exclusion mechanism. Interestingly, we observed changes to the cell morphology of infected UNSW_1.7 in the late stage of infection, even though the respective cells do not lyse (Figure S19). We assume that the HRTV-DL1 mechanism responsible for cell lysis is partially active on integrated HRTV-DL1_1.7 upon HRTV-DL1 infection, however, lysis does not occur. ORF26, the predicted glycosyl hydrolase, could be responsible for de-glycosylation and destabilization of the S-layer, and thereby for changes of the cell morphology. This destabilization could subsequently cause the reduced adsorption of HRTV-DL1 (Figure S18). This also hints towards ORF6 as a holin, because this holin candidate gene is lost in HRTV-DL_1.7. However, it remains to be elucidated which mechanisms interferes with HRTV-DL1 genome replication, virion production and cell lysis.

#### *Escape mutant ACAM34_UNSW_2.14 is fully resistant to HRTV-DL1 infection, experienced a large deletion and hints towards the S-layer gene as HRTV-DL1 receptor*

Adsorption assays revealed that HRTV-DL1 does not bind to UNSW_2.14 in detectable amounts (Figure S17). No lysis was observed in infected cultures of UNSW_2.14 (Figure S18), both control and infected cultures grow slightly slower than the parental strain, and the virus genome could not be detected in cells by PCR. Virus particles are not produced by infected UNSW_2.14 (Table S14) and no reads were detected for the virus in sequencing data of UNSW_2.14. In conclusion, HRTV-DL1 is not integrated and UNSW_1.7 is fully resistant to HRTV-DL1 infection.

Genome analyses of UNSW_2.14 revealed a large gap covering the entire integrated CHR2 (Figure S22) with two interruptions, indicating a massive rearrangement in several independent events. The region between ACAM34_UNSW_01838 (Hlac_2826) and UNSW_01867 (Hlac_2855), both transposases, is still present, as well as the region between the two transposases UNSW_01829 (Hlac_2819) and UNSW_01802 (transposase inserted into Hlac_2793). The coverage of these two regions is identical with the remaining chromosome, we therefore conclude that they are still integrated. We detect a few other changes, including twelve SNPs in intergenic regions, two silent SNPs in ORFs, three SNPs that lead to a aa substitution and one insertion of two aa. Two of the aa substitutions were already detected in UNSW_1.7, additionally, we observed an aa substitution (valine to glutamic acid) in the middle of UNSW_01036, a predicted methylmalonyl-CoA mutase (Table S13). Surprisingly, UNSW_00427 (Hlac_0412), one of the two S-layer proteins (Hlac_2976 and Hlac_0412) in ACAM34, experienced a deletion of two aa (D45 and S46) in the C-terminus. Additionally, the second S-layer protein, Hlac_2976, is removed by the large deletion. Since adsorption of HRTV-DL1 to UNSW_2.14 is fully abolished, we conclude that the S-layer represents the receptor for HRTV-DL1. Variability of S-layer genes of ACAM34 was previously detected by two ‘omics’-based studies of Deep Lake, and suggested to be driven by an arms race between viruses and host [2, 3]. Our results strongly support this hypothesis. We also propose that the deletion of Hlac_2976 causes the slightly reduced growth rate of UNSW_2.14. Cell morphology, as observed by light microscopy, does not seem to be significantly impacted (Figure S23), suggesting that Hlac_0421 alone is able to maintain the S-layer. However, determining detailed structural differences and thereby maybe differences in the stability of the S-layer comparing ACAM34_UNSW, expressing both S-layer proteins, and UNSW_1.7, with only one S-layer protein, will require high resolution imaging [4].

#### *Escape mutant ACAM34_UNSW_3.3 is partially resistant to HRTV-DL1 infection and exhibits an insertion into the S-layer gene and the deletion of an origin of replication*

A weak binding of HRTV-DL1 to UNSW_3.3 was detected by adsorption assays (Figure S17) and the virus genomes was detected in infected cells. However, production of virus particles was not observed (Table S14) and lysis was also not detectable by optical density measurements (Figure S18). Interestingly, both control and infected cultures grow slightly faster than the parental strain. In conclusion, UNSW_3.3 is partially resistant to HRTV-DL1 adsorption and is able to prevent cell lysis, even though the virus genome is to some extend replicated in host cells.

Genome sequencing of UNSW_3.3 recovered no reads for the virus genome, but revealed a number of mutations on the chromosome (Table S13). We found 12 SNPs in intergenic regions, the two aa substitutions observed in the other two escape mutants, the frame shift in ACAM34UNSW_01960 that was also detected in UNSW_1.7 and two silent SNPs. We do not expect that any of these mutations has an influence on the virus life cycle, since the majority of them are also present in other escape mutants that have different phenotype. The major change influencing virus-host interactions is an insertion of 12 aa (TPPTVSRLCFDT) between aa209 and aa210 of ACAM34UNSW_01982 (Hlac_2976), the S-layer protein that is also affected in UNSW_2.14. However, the mutation only accounts for (90 %) of the population, suggesting that the drastically reduced adsorption is caused by this mutation and some virus particles can adsorb to the remaining 10 % of the population. Hlac_0412 did not experience a mutation in UNSW_3.3, but we still see a dramatically reduced adsorption, indicating that either Hlac_2976 is the preferred receptor of HRTV-DL1, or the Hlac_2976 is preferably used by the cell to build the S-layer. Nevertheless, this does not explain why we do not detect virus particles in the supernatant of the infected culture considering that 10 % of the population can be infected. Another major change in UNSW_3.3 is the deletion of 66 nt upstream of UNSW_01845 (Hlac_2833 CHR2), a gene coding for cdc6/orc1. This deletion affects an origin of replication (ORI) that is most likely recognised by the adjacent cdc6 and explains the slightly enhanced growth rate of the escape mutant. Deletions of ORIs in *Haloferax volcanii* were shown to provide a growth advantage [5]. One other escape mutant, ACAM34_UNSW_2.10, that was not further analysed showed a similar phenotype and might exhibit the same mutation. The only other significant change are two substitutions (M->V) in ACAM34UNSW_02839 (Hlac_2475), however, this change is only found in 25 % of the population and no function could be predicted for the respective ORF.

It is difficult to hypothesize how the deletion of an ORI, the binding site for the host origin recognition protein (CDC6/ORC1), could prevent production of virus particles. However, the deletion of an ORI could redirect the CDC6 protein from the ORI to another target, which is likely more effective than the differential regulation of the gene. Indeed, we also do detect differential expression of orc1/cdc6 genes under viral infection. While we only detected one of fifteen annotated orc1/cdc6 genes (Hlac_3217) being upregulated in resistant ACAM34_DSMZ, five of seven remaining orc1/cdc6 genes are differentially regulated in sensitive ACAM34_UNSW (ACAM34UNSW_01140 is upregulated at T1 and downregulated in T2; ACAM34UNSW_01549, ACAM34UNSW_01845, ACAM34UNSW_01965 and ACAM34UNSW_01106 are upregulated at T2) (Table S8), including the orc1/cdc6 gene adjacent to the destroyed origin. Upregulation of orc1/cdc6 genes is in general rather unexpected, given that viruses tend to reprogram the cell towards replication of the virus genome and some viruses even inhibit host genome replication to have the pool of nucleotides available for virus replication [6]. This indicates that upregulation of orc1/cdc6 genes is an active host response to viral infection. We could not identify any sequences similar to the deleted ORI on the HRTV-DL1 genome that could represent an alternative binding site for CDC6 proteins. However, studies have shown that CDC6 can bind dsDNA without sequence specificity [7], that dysregulation of CDC6 expression can lead to inhibition of replication [8] and that CDC6 is also able to recruit the RNA polymerase I for rDNA transcription initiation [9]. Therefore, we suggest that the overrepresentation of CDC6 proteins in infected host cells could be involved in inhibiting virus replication or even transcription and thereby inhibit assembly of virus particles. Alternatively, ATP depletion has been shown to be a strategy of virus exclusion, which could be facilitated by CDC6 protein [10], CDC6 proteins are ATPases. Nevertheless, the potential anti-viral activity of CDC6 in ACAM34 needs to be investigated experimentally.

### Supplementary Figures

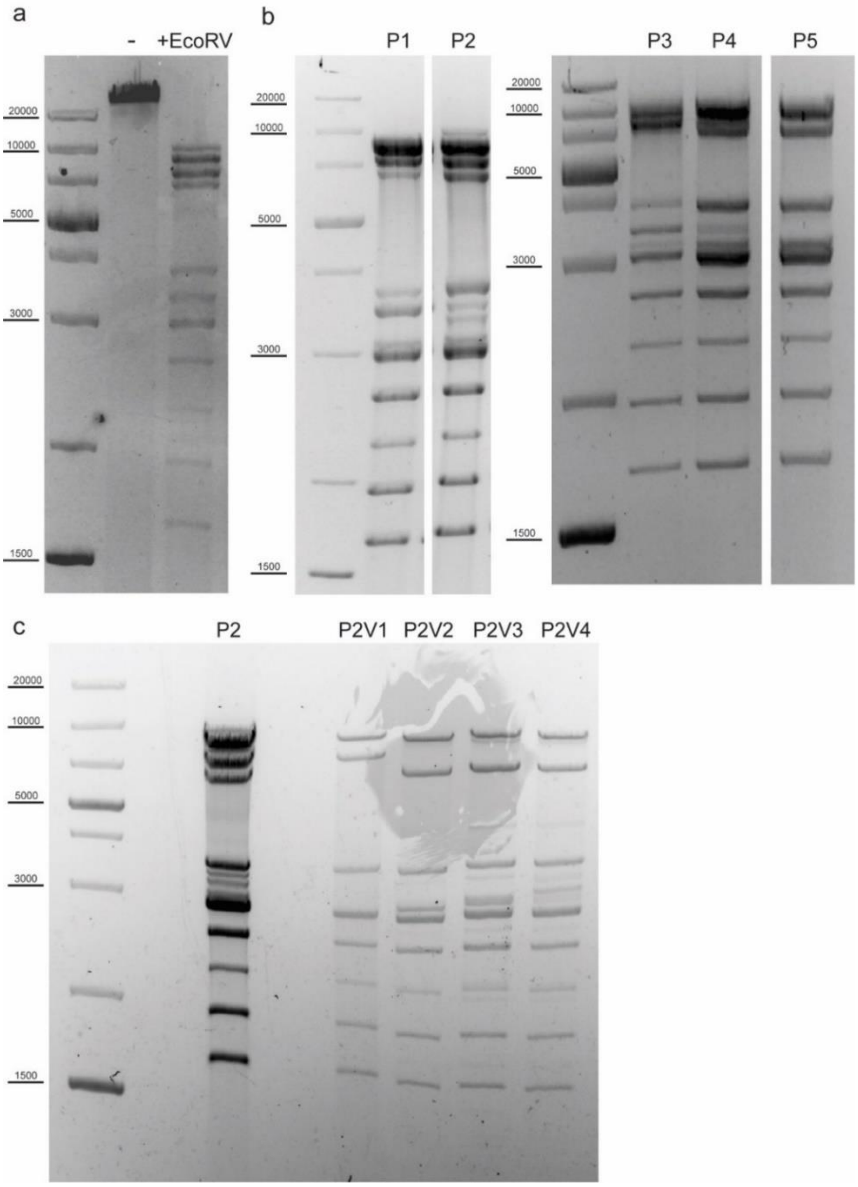


**Figure S1. Analysis of genomic DNA of the original virus isolate and different variants isolated from single plaques.** (**a**) Total DNA from virus particles isolated from the supernatant of a *Hrr. lacusprofundi* isolate [2] (HRTV-DL) undigested (-) and digested with *Eco*RV (+EcoRV). (**b**) EcoRV digest of DNA from HRTV-DL variants isolated from single plaques (P1-P5), formed after infection with the original virus preparation on *Hrr. lacusprofundi* ACAM34 lawns and propagated in the same strain. (**c**) EcoRV digest of DNA from HRTV-DL variants isolated from single plaques (P2V1-V4) formed after infection with P2 (same as in b) on *Hrr. lacusprofundi* ACAM34 lawns and propagated in the same strain. P2V1 (HRTV-DL1) was chosen for further studies. MW size marker is shown to the left of the gel (GeneRuler 1 kb Plus DNA Ladder, Thermo Fisher Scientific). DNA was separated on 1% agarose gels and stained with SYBR™ Safe DNA stain. Original gel images have been modified by excising separated lanes to improve visual presentation.

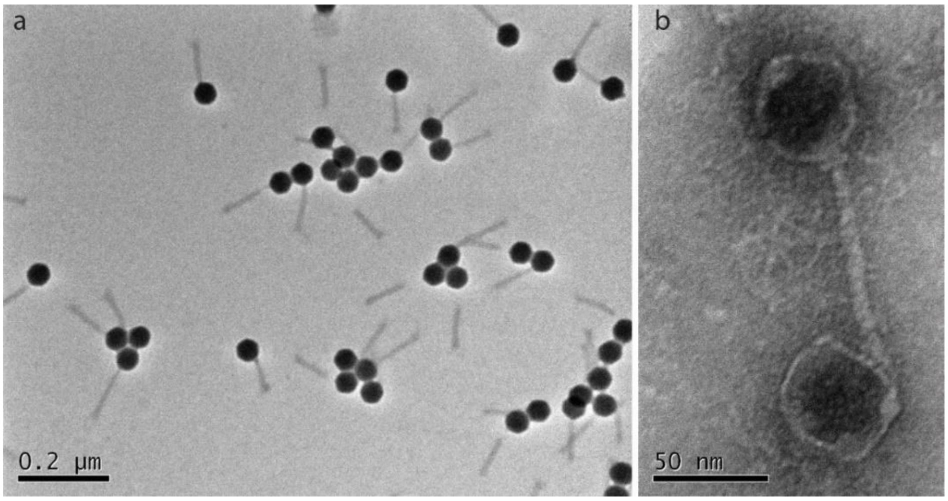


**Figure S2. Transmission electron micrographs HRTV-DL1.** (**a**) HRTV-DL1 virus particles purified from *Hrr. lacusprofundi* ACAM34 cultures. (**b**) Detailed view of HRTV-DL1 virus particle attached to a membrane vesicle. Samples were negatively stained with 2% uranyl acetate.

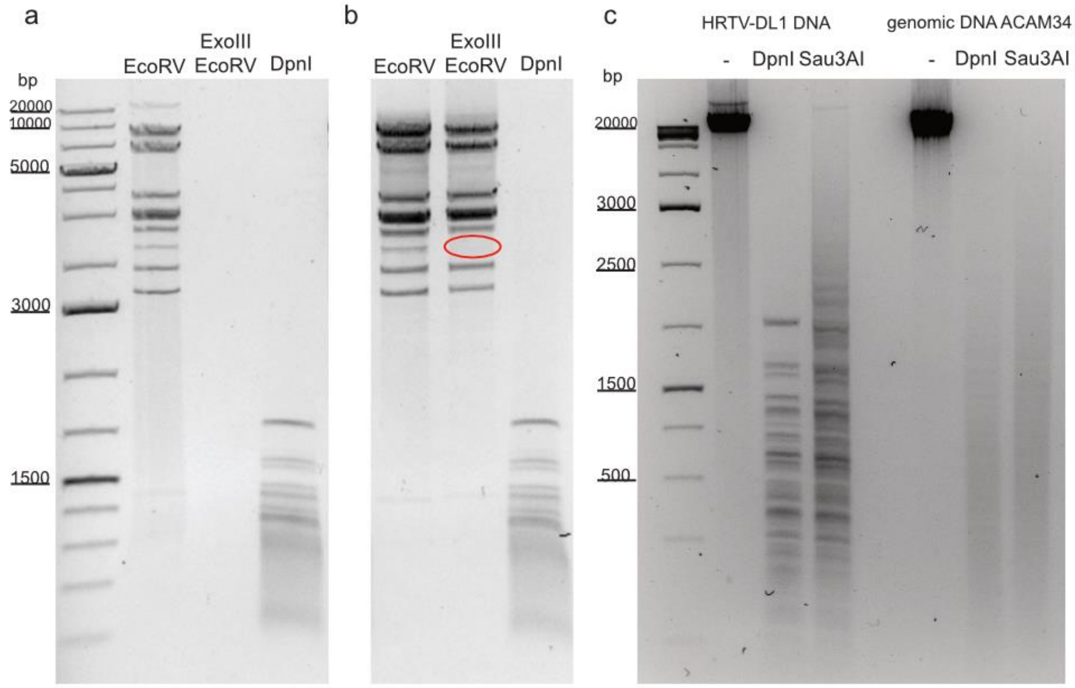


**Figure S3. Analysis of genomic DNA of HRTV-DL1.** (**a**) Total DNA of HRTV-DL1 isolated from purified virus particles digested with *Eco*RV, with Exonuclease III and subsequently *Eco*RV, and with *Dpn*I. (**b**) HRTV-DL1 DNA isolated from infected host cells (*Hrr. lacusprofundi* ACAM34) digested with *Eco*RV, with Exonuclease III and subsequently *Eco*RV, and with *Dpn*I. A band that is not present after Exonulcease III digest, is marked with a red circle. (**c**) Total DNA of HRTV-DL1 and *Hrr. lacusprofundi* ACAM34 undigested (-) and digested with *Dpn*I and *Sau*3AI. MW size marker is shown to the left of the gel (GeneRuler 1 kb Plus DNA Ladder, Thermo Fisher Scientific). DNA was separated on 1% (a,b) and 2% (c) agarose gels and stained with SYBR™ Safe DNA stain. Original gel images have been modified by cropping to improve visual presentation.

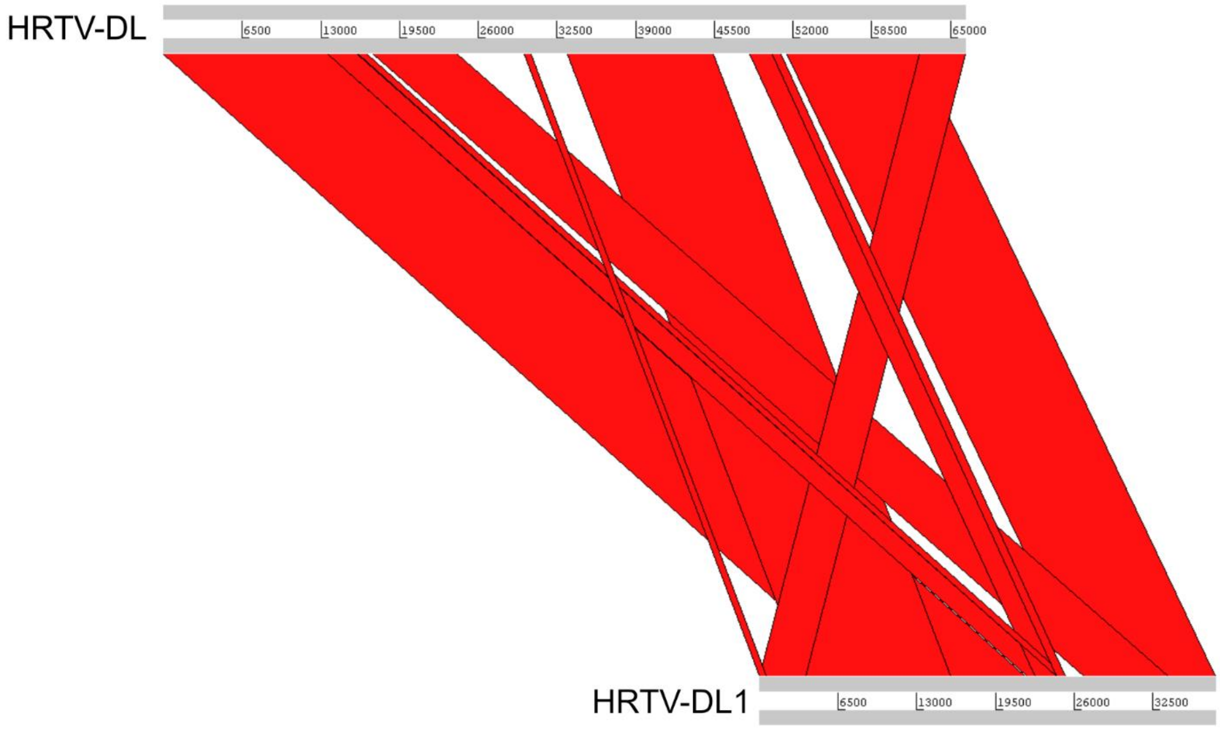


**Figure S4. Genome comparison of HRTV-DL and HRTV-DL1.** Nucleotide sequence of genomes was compared using Basic Local Alignment Search Tool (BlastN) and comparison visualized using Artemis comparison tool (ACT)[11].

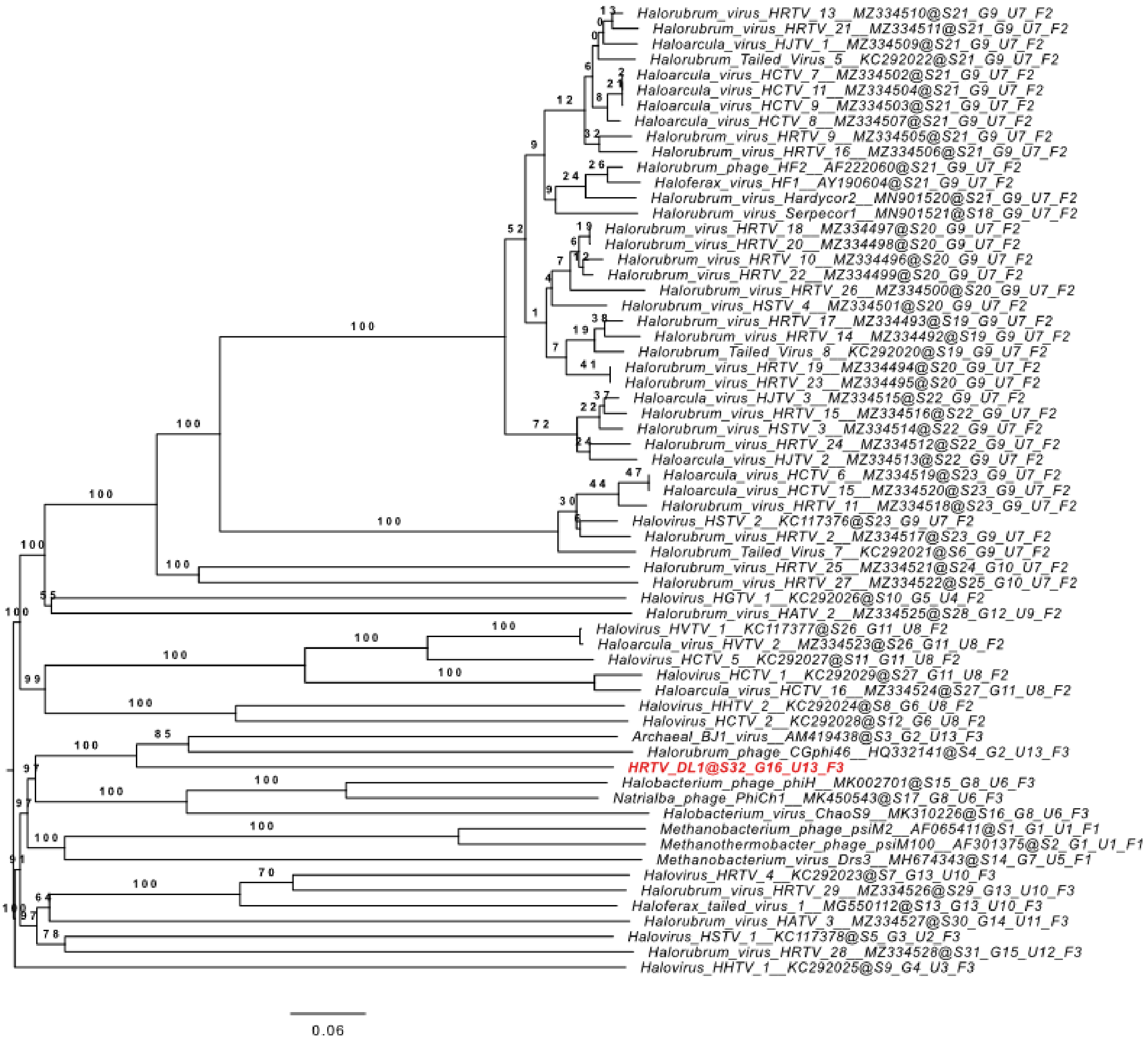


**Figure S5. Phylogenetic tree** reconstructions from protein sequences of 63 arTVs and HRTV-DL1 (highlighted in red). Phylogenomic GBDP trees inferred using the formulas d6 and yielding average support of 54 %. The numbers above branches are GBDP pseudo-bootstrap support values from 100 replications. The branch lengths of the resulting VICTOR trees are scaled in terms of the respective distance formula used.

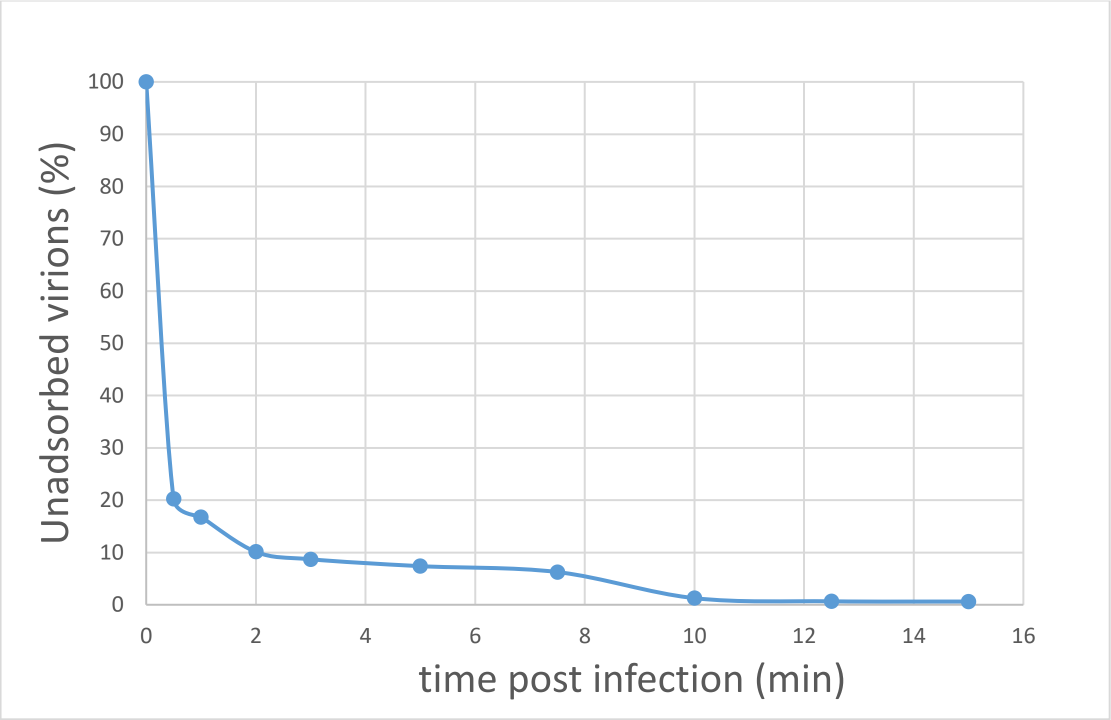


**Figure S6. Adsorption of HRTV-DL1 to cells of *Hrr. lacusprofundi* ACAM34_UNSW**. Cells were infected with HRTV-DL1 at room temperature (20°C). The number of unbound virus particles was determined at different time points post infection by plaque assay. Graph represents one of three biological replicates.

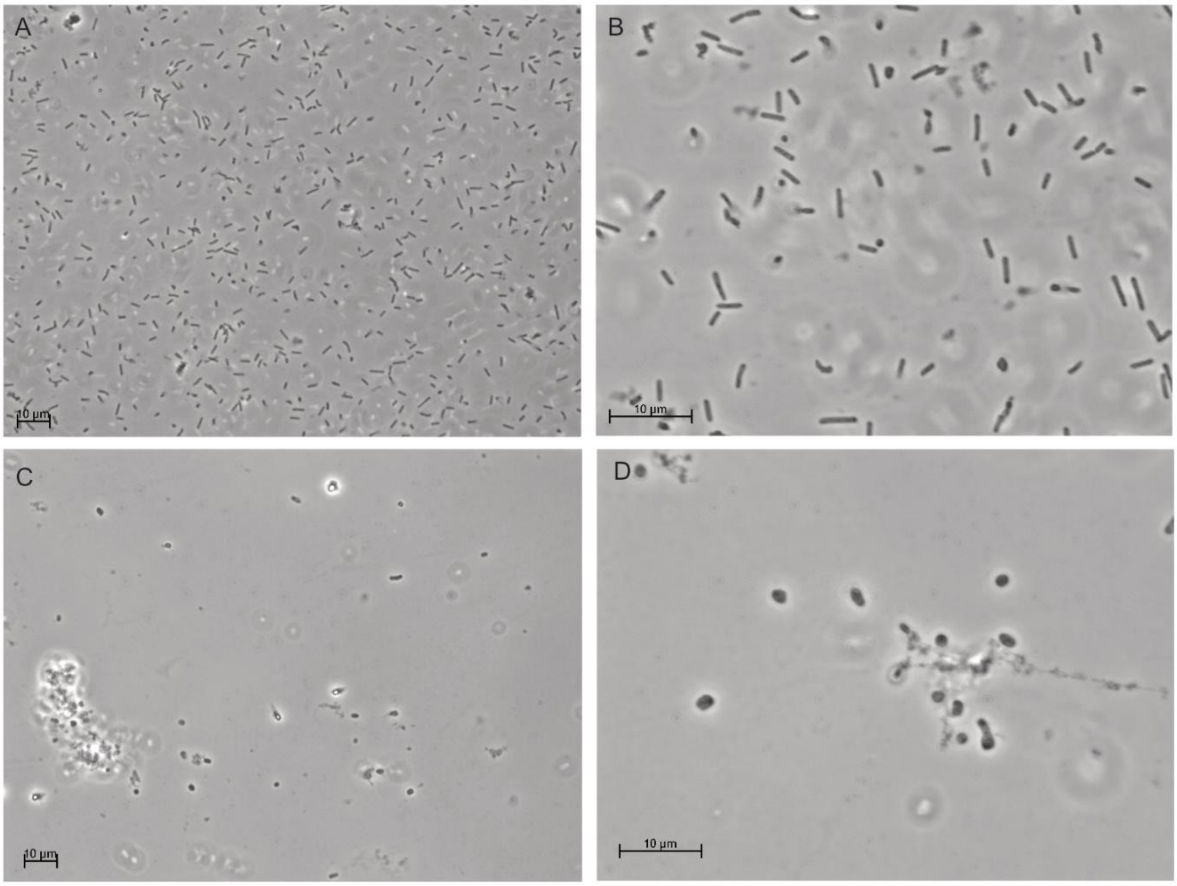


**Figure S7. Cell-shape changes of HRTV-DL1 infected *Hrr. lacusprofundi* ACAM34_UNSW**. Light Microscopy of uninfected (A and B) and HRTV-DL1 infected (C and D) *Hrr. lacusprofundi* 42 hours post infection at the onset of cell lysis. Cells are fixed with 1% glutaraldehyde.

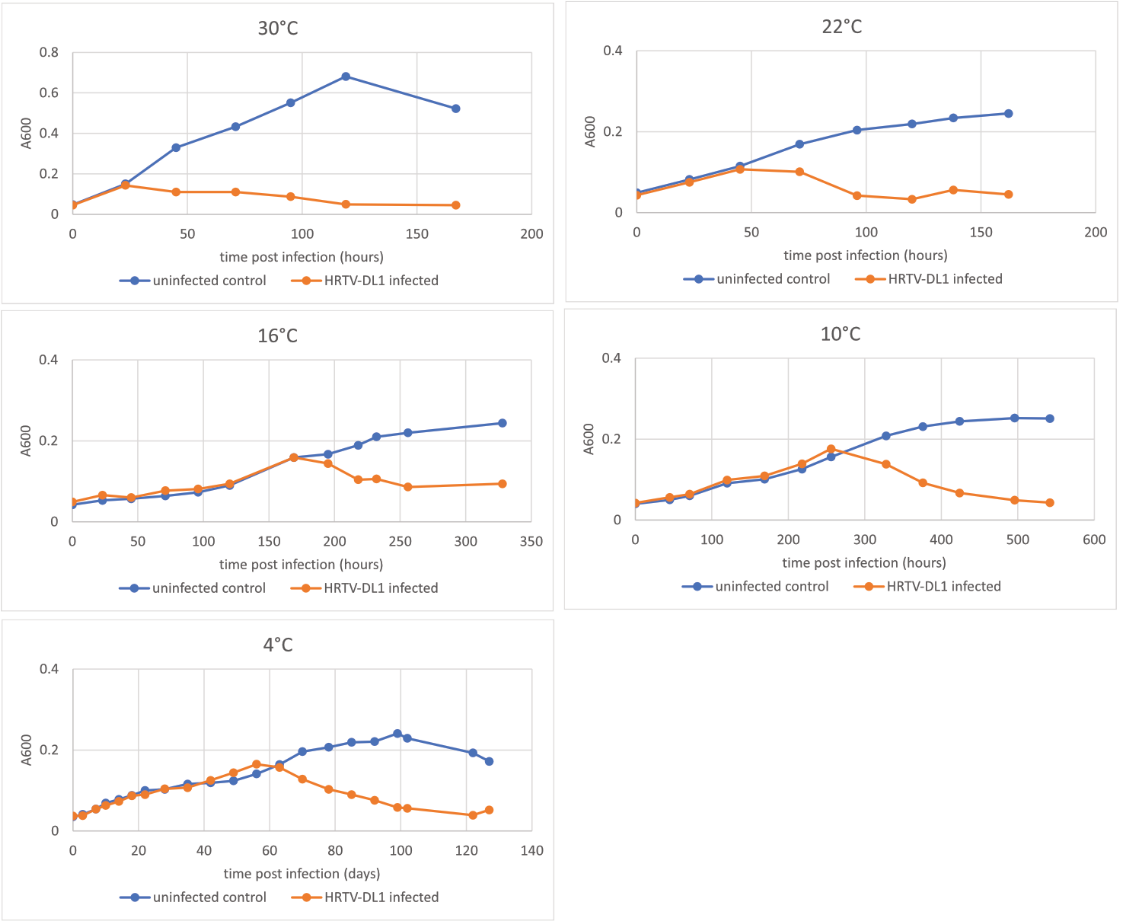


**Figure S8. Virus life cycle at different temperatures.** Growth curves of uninfected control and HRTV-DL1 infected *Hrr. lacusprofundi* ACAM34_UNSW at different temperatures.

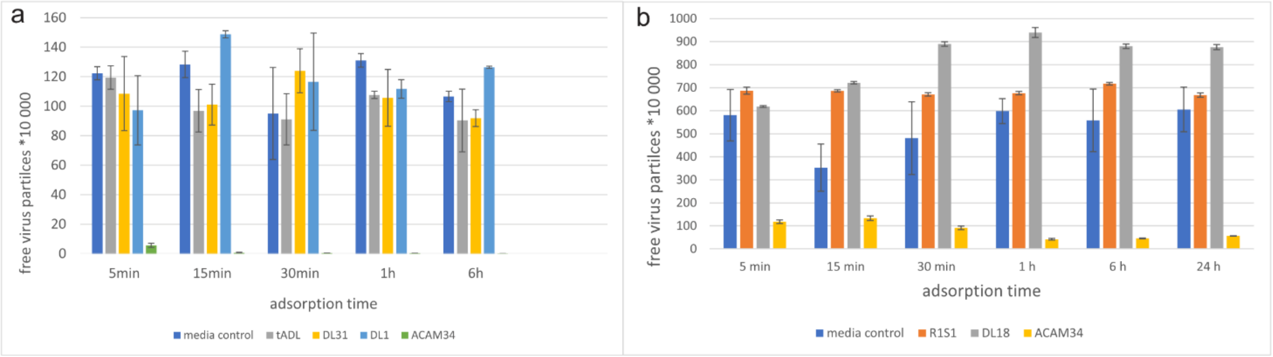


**Figure S9. Adsorption assay of HRTV-DL1 with different potential host organisms.** (**a**) *Halohasta litchfieldiae* tADL, *Hrr. lacusprofundi* ACAM34_DSMZ, halophilic archaeon DL31, and *Halobacterium* DL1. (**b**) *Hrr. lacusprofundi* ACAM34_DSMZ, R1S1 and DL18. Virus particles were incubated with host cells (or cell free media as control) for 5,15,30,60 and 360 min and number of not-adsorbed free virus particles in the supernatant was determined by plaque assay. Graphs represent one of three biological replicates. Error bars represent standard deviation from three independent experiments.

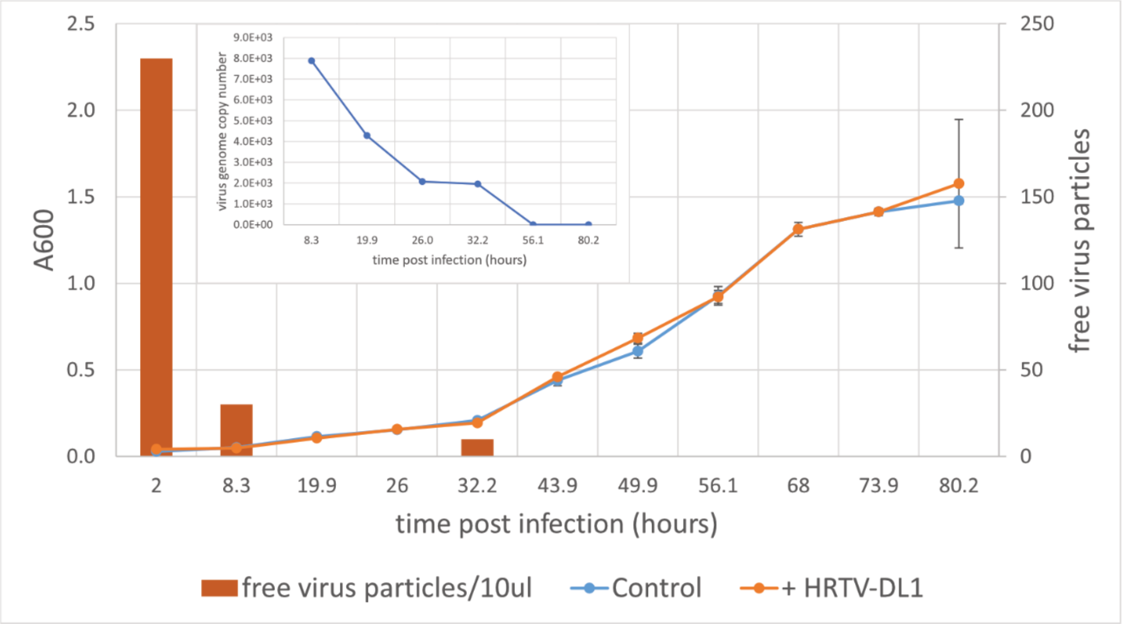


**Figure S10. Infection of *Hrr. lacusprofundi* ACAM34_DSMZ with HRTV-DL1.** Growth curve of uninfected control and HRTV-DL1 infected *Hrr. lacusprofundi* ACAM34_DSMZ. Free virus particles in 10ul culture supernatant were determined by plaque assay. **Inlet:** Virus genome copy number determined by per ml cell culture by qPCR. Samples were taken from the growth curve shown in the main figure. Samples in which virus genomes could not be detected by qPCR were set to 0. Graphs represent one of three biological replicates. Error bars represent standard deviation from three independent experiments.

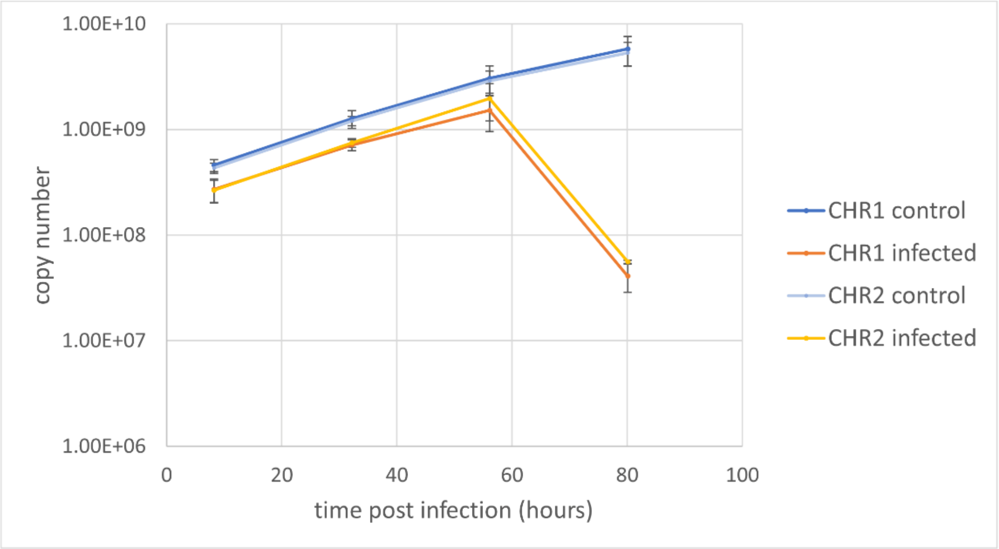


**Figure S11. Copy numbers of ACAM34_UNSW main chromosome (CHR1) and the integrated secondary chromosome (CHR2)** in uninfected controls and HRTV-DL1 infected cultures. Copy numbers were determined from 2ml cell culture by qPCR and samples were taken from the growth curve shown in Figure 5. Graphs shown are averages of three biological replicates. Error bars represent standard deviation from three independent experiments.

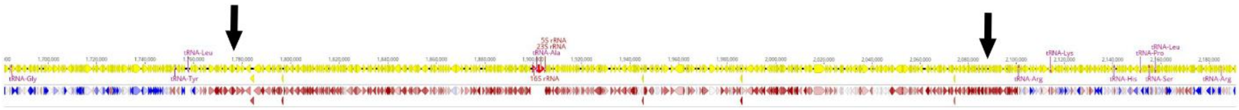

**Figure S12. Upregulation of former CHR2 in ACAM34_UNSW under viral infection.** Section of the genome of ACAM34_UNSW with CHR2 integrated and expression level of genes compared between uninfected control samples and infected samples. The level of differential expression is shown in the lower panel with different shades of blue representing different levels of downregulation and different shades of red representing different levels of upregulation. Arrows indicate the borders of former CHR2. Visualized using Geneious version 2022.1 created by Biomatters (https://www.geneious.com).

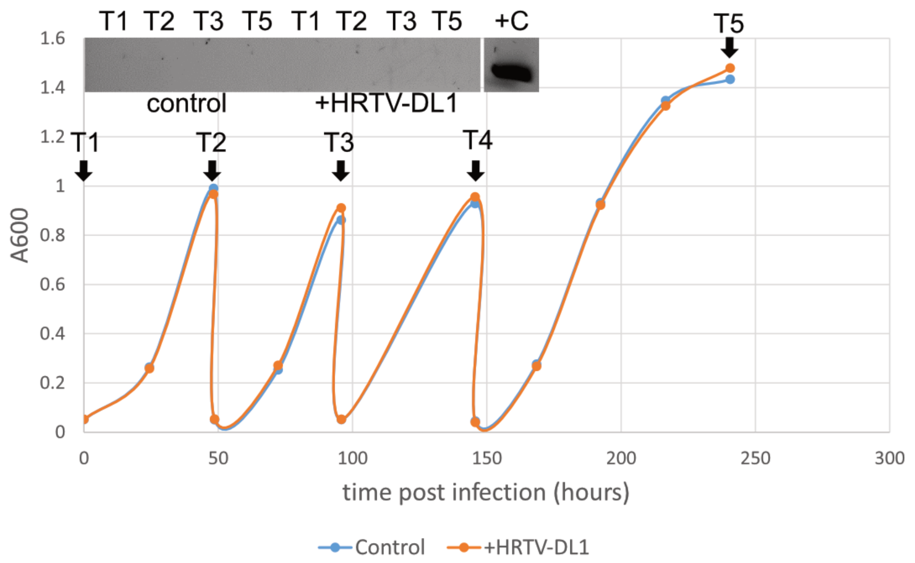


**Figure S13. Long-term infection of *Hrr. lacusprofundi* ACAM34_DSMZ with HRTV-DL1.** Growth curve of uninfected control and HRTV-DL1 infected *Hrr. lacusprofundi* ACAM34_DSMZ, with serial dilution of the culture in exponential growth upon reaching A600nm=1 (T2-T3). **Inlet:** PCR on HRTV-DL1 in ACAM34_DSMZ cells at indicated times (T1, T2, T4 and T5) post infection. +C positive control (HRTV-DL1 DNA). Graphs represent one of three biological replicates. Original gel images have been modified by excising separated lanes to improve visual presentation. DNA was separated on 1% agarose gels and stained with SYBR™ Safe DNA stain.

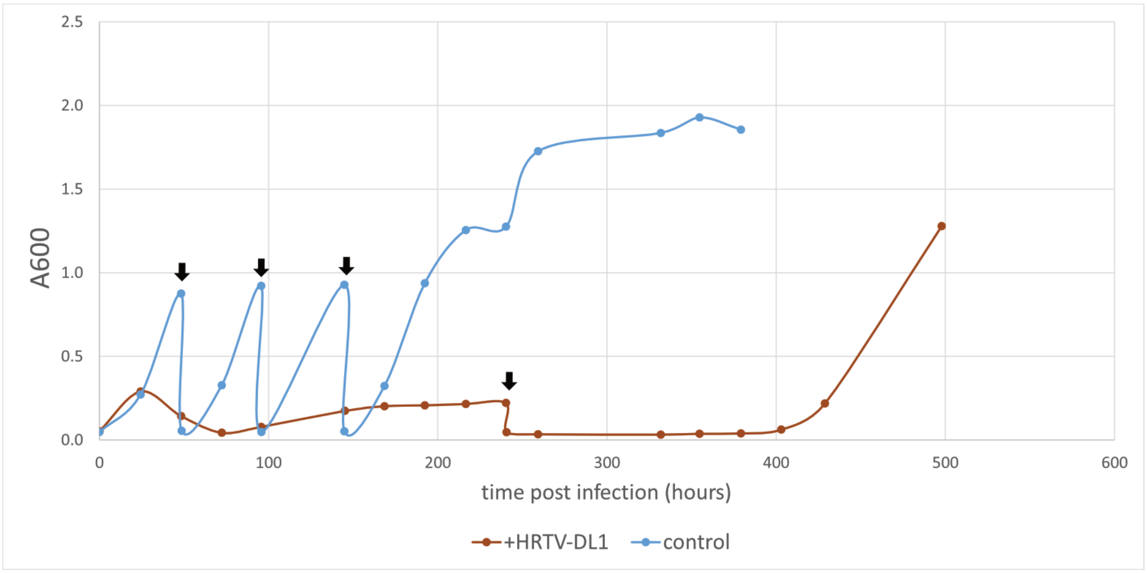


**Figure S14. Long-term infection of *Hrr. lacusprofundi* ACAM34_UNSW with HRTV-DL1.** Growth curve of uninfected control and HRTV-DL1 infected *Hrr. lacusprofundi* ACAM34_UNSW, with serial dilution of the culture in exponential growth as indicated by black arrows. Graphs represent one of three biological replicates.

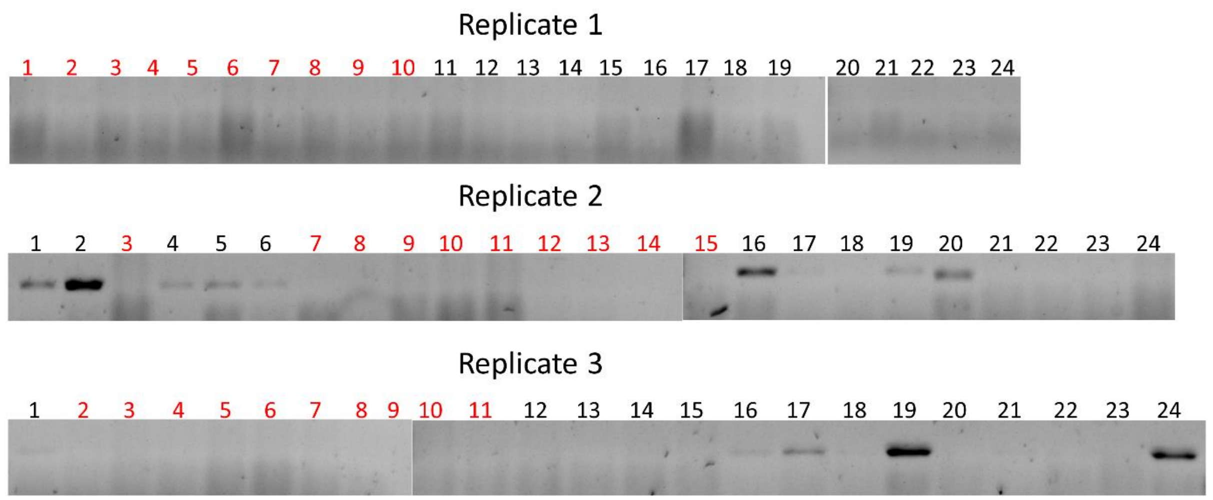


**Figure S15. HRTV-DL1 in single colonies obtained from long-term infection of *Hrr. lacusprofundi* ACAM34_UNSW with HRTV-DL1.** PCR on HRTV-DL1 in ACAM34_UNSW single clones obtained from three biological replicates of recovered long-term infected ACAM34_UNSW cultures. Colonies labeled with red numbers were chosen for further characterization. Original gel images have been modified by excising separated lanes to improve visual presentation. DNA was separated on 1% agarose gels and stained SYBR™ safe DNA stain.

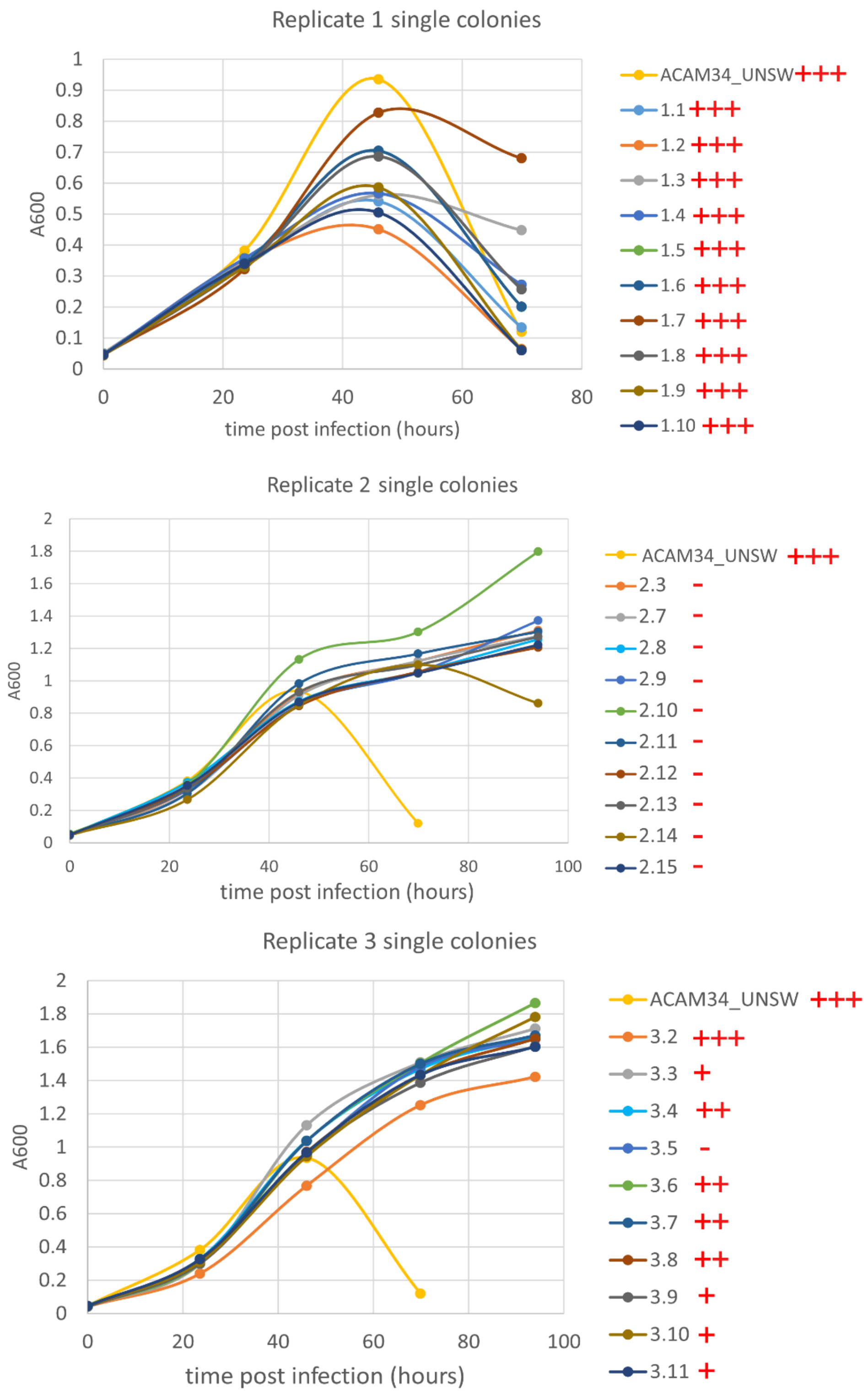


**Figure S16: Infection of *Hrr. lacusprofundi* ACAM34_UNSW single colonies retrieved from virus recovered cultures.** Growth curve of HRTV-DL1 infected *Hrr. lacusprofundi* ACAM34_UNSW (as control) and re-infected virus free single colonies retrieved from recovered HRTV-DL1 infected cultures (10 colonies per biological replicate). Infection with HRTV-DL1 was detected by PCR and is indicated with +++ for a very strong signal, ++ for a strong signal, + for a weak signal and – for no signal.

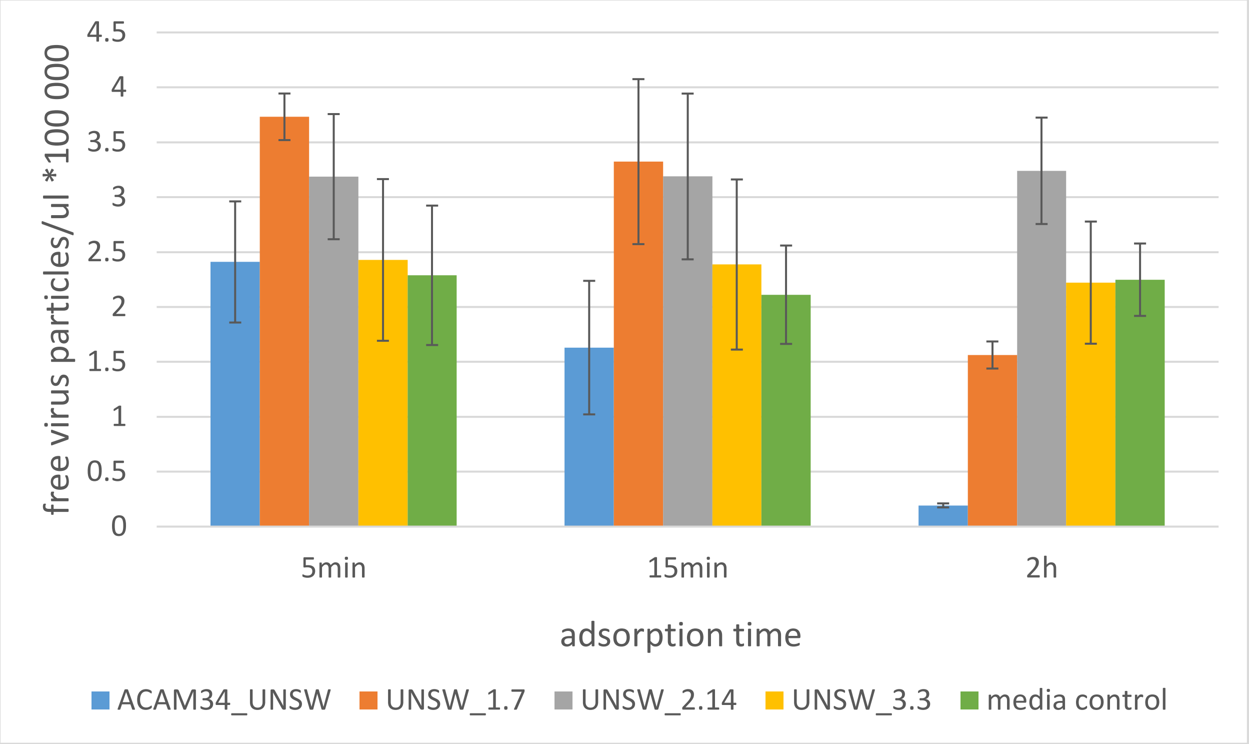


**Figure S17. Adsorption assay of HRTV-DL1 with strains recovered from HRTV-DL1 infected ACAM34_UNSW cultures.** Virus particles were incubated with host cells (or cell free media as control) for 5 min, 15min and 2 hours and number of non-adsorbed free virus particles in the supernatant was determined by plaque assay. Graphs represent average values of three biological replicates. Error bars represent standard deviation from three independent experiments.

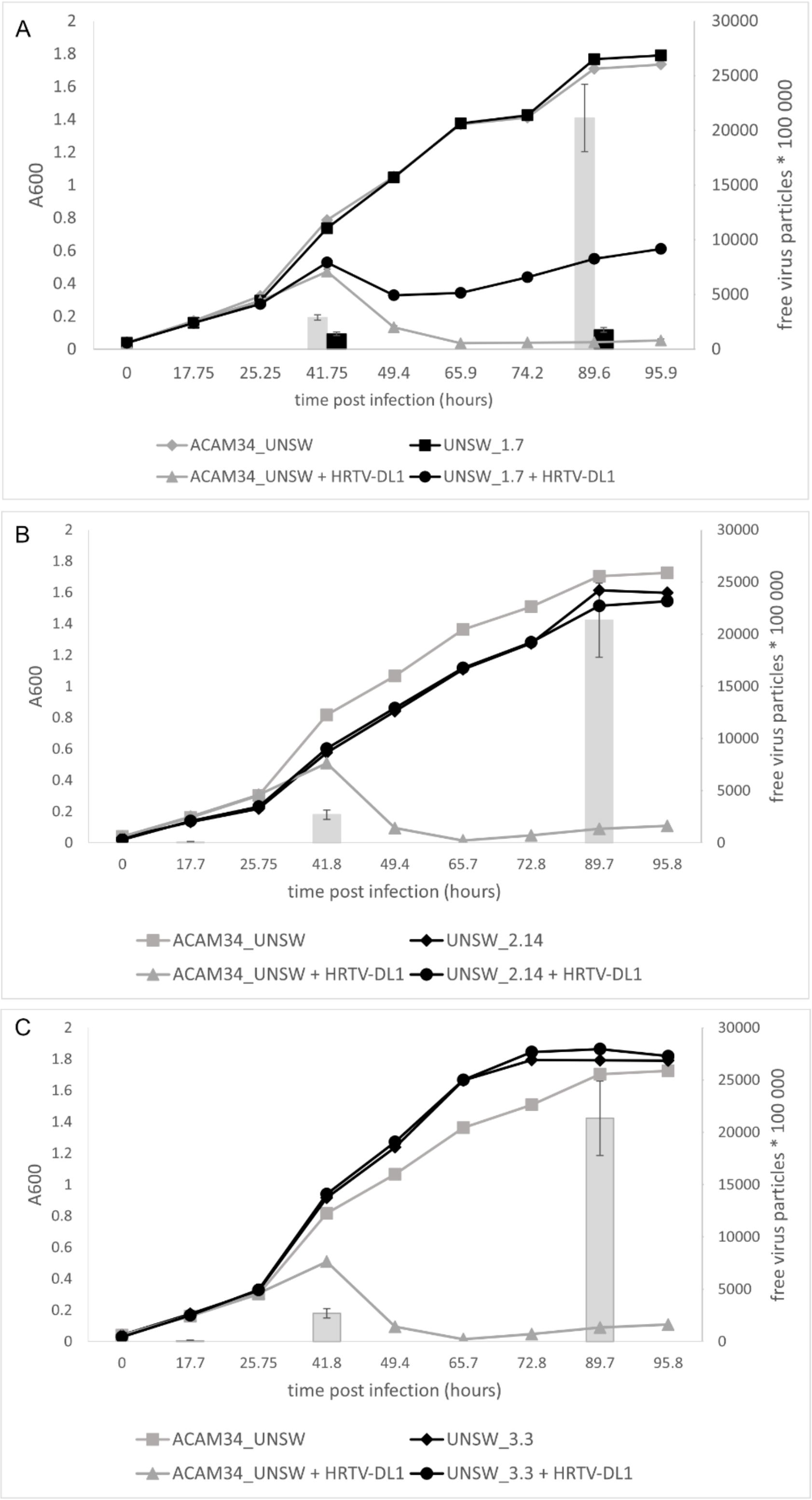


**Figure S18. Life cycle of HRTV-DL1 in ACAM34_UNSW escape mutants**. Growth curves of uninfected control and HRTV-DL1 infected *Hrr. lacusprofundi* ACAM34_UNSW and escape mutants UNSW_1.7 (A), UNSW_2.14 (B) and UNSW_3.3 (C). Free virus particles in 10ul culture supernatant were determined by plaque assay and are presented by grey bars (parental strain) and black bars (escape mutants). Graphs represent one of three biological replicates.

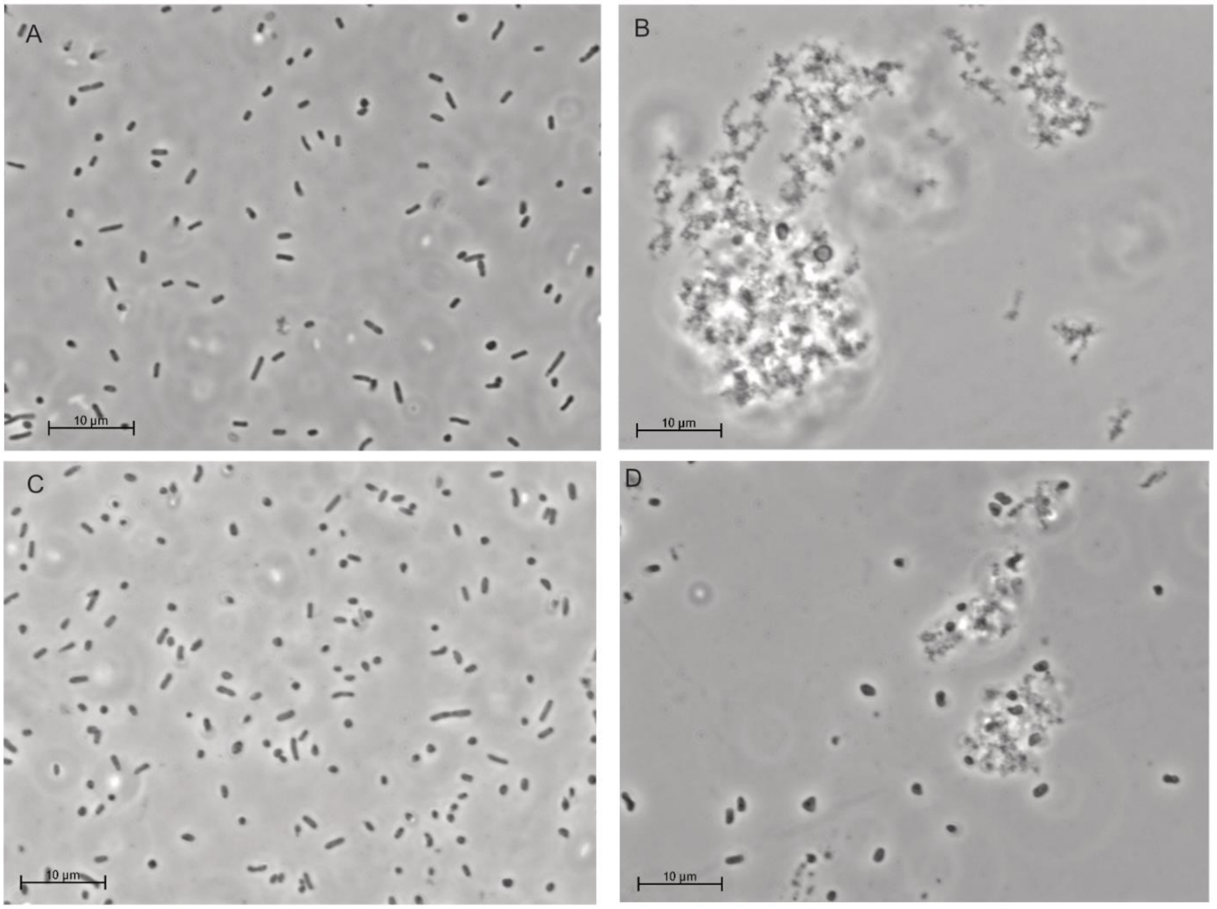


**Figure S19. Cell-shape changes of HRTV-DL1 infected *Hrr. lacusprofundi* UNSW_1.7**. Light Microscopy of uninfected (A) and HRTV-DL1 infected (B) ACAM34_UNSW parental strain in comparison with uninfected (C) and HRTV-DL1 infected (D) UNSW_1.7 at 90 hours post infection.

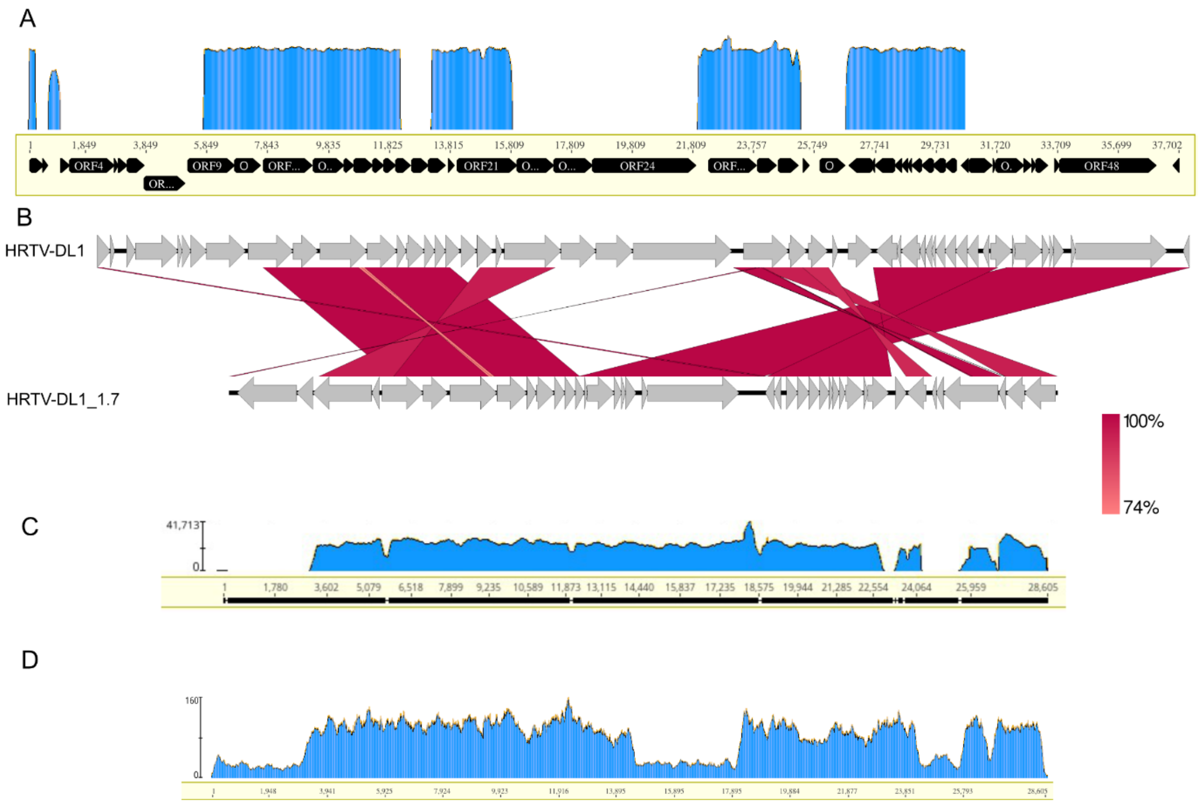


**Figure S20. Genome of HRTV-DL1_1.7. (A)** Read mapping of UNSW_1.7 genome sequencing data onto the HRTV-DL1 genome. Visualized using Geneious version 2022.1 created by Biomatters. **(B)** Genome comparison of HRTV-DL1_1.7 with HRTV-DL1 using easyfig [12]. **(C)** Read mapping of HRTV-DL1 genome sequencing data onto the HRTV-DL1_1.7 genome. **(D)** Read mapping of HRTV-DL genome sequencing data onto the HRTV-DL1_1.7 genome. Visualized using Geneious version 2022.1 created by Biomatters (www.geneious.com).

.

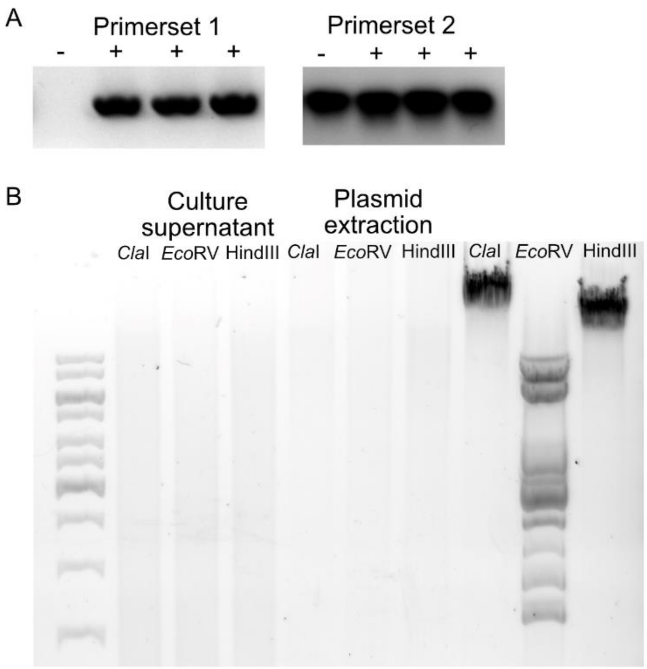


**Figure S21. Detection of HRTV-DL1_1.7. (A)** PCR detection of HRTV-DL1_1.7 in UNSW_1.7 with primers (Primerset 1) used to detect HRTV-DL1 (HRTV-DLF and HRTV-DLR, Supplementary Table 1) did not detect HRTV-DL1_1.7 in UNSW_1.7. Primers (Primerset 2) used to determine copy numbers of HRTV-DL1 by qPCR (BV37VPF and BV37VPR, Supplementatry Table 1) detect HRTV-DL1_1.7 in UNSW_1.7. (-) uninfected UNSW_1.7, (+) UNSW_1.7 infected with HRTV-DL1. Original gel image has been modified by excising separated lanes to improve visual presentation. **(B)** Restriction digest of DNA isolated from culture supernatants (lanes 2-4) and plasmid preparation (lanes 5-6) of cells of UNSW_1.7 after UV treatment in comparison with HRTV-DL1 infected ACAM34_UNSW wild type (lane 7-9). MW size marker is shown to the left of the gel (GeneRuler 1 kb Plus DNA Ladder, Thermo Fisher Scientific). DNA was separated on 1% agarose gels and stained with SYBR™ safe DNA stain.

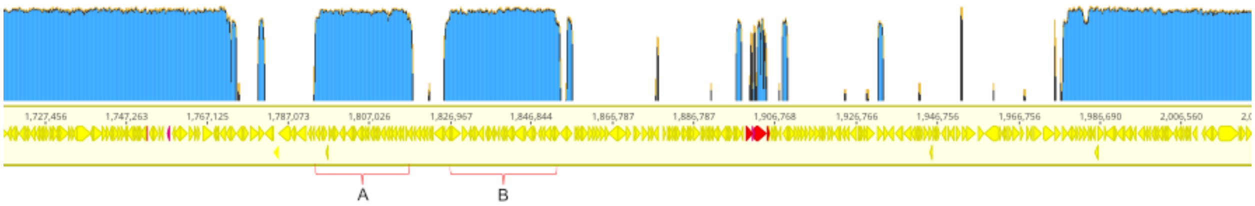


**Figure S22. Genome of escape mutant ACAM34_UNSW_2.14.** Read mapping of UNSW_2.14 genome sequencing data onto the ACAM34_UNSW genome (log scale). Only the section with the deletion of CHR2 is shown. A and B are sections of CHR2 that are enclosed between two transposases and are still present. Remaining regions with coverage within the gap are either transposases and ribosomal RNA that recruit reads due to their high sequence similarity, or intergenic regions. Visualized using Geneious version 2022.1 created by Biomatters (www.geneious.com).

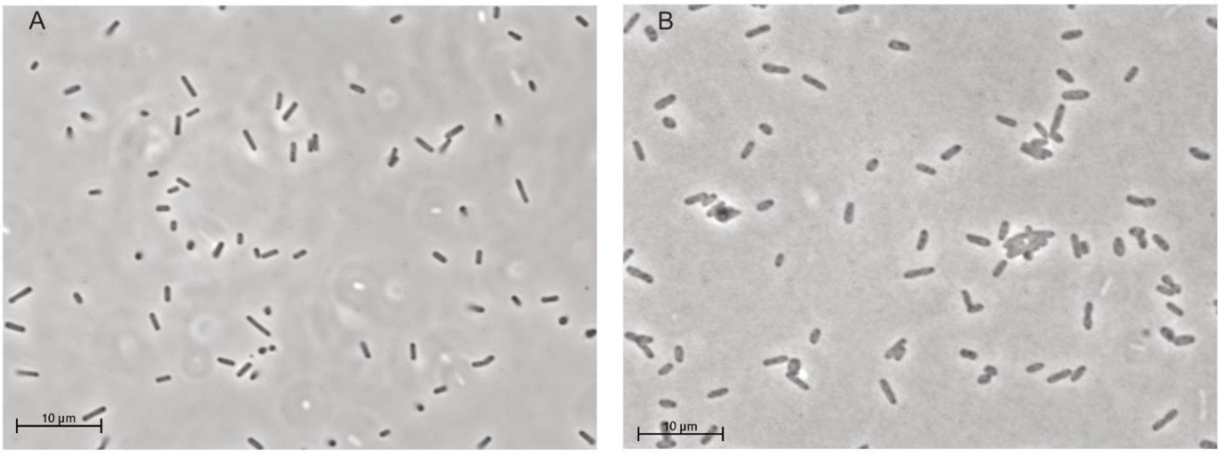


**Figure S23. Cell morphology of *Hrr. lacusprofundi* UNSW_2.14**. Light microscopic images of (A) ACAM34_UNSW parental strain in comparison with (B) UNSW_2.14.

### Supplementary Tables

**Table S1. Blast result comparing annotated ORFs from HRTV-DL with HRTV-DL1.** (Excel file)

**Table S2. Annotation of HRTV-DL1 genome and identification of proteins in the virus particle.** (Excel file)

**Table S3. Proteins identified in purified virus particles.** (Excel file)

**Table S4. CRISPR spacer hits of Deep Lake haloarchaea against HRTV-DL1.** (Excel file)

**Table S5. Putative virus defense mechanisms in ACAM34_DSMZ.** Output from PADLOC webserver [13] run on ACAM34_DSMZ. (Excel file)

**Table S6. Differential expression of ACAM34_DSMZ genes under infection with HRTV-DL1.** Differential expression at time point 1 (56 hours post infection), calculated with Geneious Prime® 2022.2.1, from 2 replicates uninfected and 3 replicates HRTV-DL1 infected. (Excel file)

**Table S7. ACAM34_DSMZ genes upregulated under HRTV-DL1 infection that are absent from sensitive strain ACAM34_UNSW.** (Excel file)

**Table S8. Differential expression of ACAM34_UNSW genes under infection with HRTV-DL1.** Differential expression at time point 1 (56 hours post infection), calculated with Geneious Prime® 2022.2.1, from 2 replicates uninfected and 3 replicates HRTV-DL1 infected. (Excel file)

**Table S9. Comparison of Expression levels of the S-layer gene (ACAM34_UNSW_01982) with Hlac_3088 (ACAM34_UNSW_02085) in ACAM34_UNSW.** Expression values (FPKM) have been calculated with Geneious Prime® 2022.2.1, and averages from 2 replicates uninfected and 3 replicates HRTV-DL1 infected are presented.

| Sample | ACAM34_UNSW_01982 | ACAM34_UNSW_02085 |
|---|---|---|
| **32 hours p.i. control** | 496 | 401 |
| **56 hours p.i. control** | 544 | 267.5 |
| **32 hours p.i. infected** | 1876 | 294 |
| **56 hours p.i. infected** | 677.33 | 849 |

**Table S10. Expression levels of HRTV-DL1 ORFs 56 hours p.i.** FPKM values for HRTV-DL1 ORFs in three different replicates, and average of those, 56.1 hours post infection. Expression values (FPKM) have been calculated with Geneious Prime® 2022.2.1. (Excel file)

**Table S11: Primer Sequences** (Excel file)

**Table S12: Open reading frames encoded in HRTV-DL1_1.7 that have not been described on the HRTV-DL1 genome.**

| ANNOTATION (LENGTH AMINO ACIDS) | NCBI HIT | CONSERVED DOMAINS |
|---|---|---|
| **ORF_00001 (684)** | hypothetical protein HrrHc1_145 [Halorubrum phage Hardycor1], 32% identity | none |
| **ORF_00002 (186)** | hypothetical protein HrrHc1_125 [Halorubrum phage Hardycor1], 32% identity | none |
| **ORF_00033 (125)** | site-specific integrase [Halorubrum saccharovorum], 56% identiy | site-specific integrase |
| **ORF_00035 (49)** | No hits | none |
| **ORF_00036 (82)** | hypothetical protein [Halobellus rufus], 49% identity | none |
| **ORF_00038 (79)** | hypothetical protein C490_03403 [Natronobacterium gregoryi SP2], 43% identity | none |

**Table S13. Variant analysis on escape mutants.** Variant analysis was performed with Geneious Prime® 2022.2.1. (Excel file)

**Table S14. Free virus particles in HRTV-DL1 infected escape mutants UNSW_2.14 and UNSW_3.3.** Free virus particles calculated for 0.0001ul culture supernatant of HRTV-DL1 infected ACAM34_UNSW and escape mutants UNSW_2.14 and UNSW_3.3, as determined by plaque assay.

| hours p. i. | ACAM34_UNSW | UNSW_2.14 | UNSW_3.3 |
|---|---|---|---|
| **41.8** | 2700 | 0.105 | 0.095 |
| **89.7** | 21345 | 0.099 | 0.052 |
